# Non-invasive brain stimulation biases temporal value–aversiveness computations and promotes sustainable decision-making

**DOI:** 10.64898/2026.02.27.708652

**Authors:** Xinyi Zhang, Wei Liu

## Abstract

Environmental sustainability depends on widespread adoption of sustainable behaviors, yet the neurocomputational mechanisms supporting such choices remain poorly understood. We developed a temporal-value decision model (TVDM) to explain sustainability-related decision-making. Behavioral data showed that sustainable decisions often carry immediate aversiveness while associated outcome values that are delayed and therefore discounted, jointly limiting sustainable decisions. We replicated these effects in an independent sample and then investigated whether these mechanisms could be externally influenced. Non-invasive transcranial direct current stimulation (tDCS) targeting the dorsolateral prefrontal cortex (DLPFC) increased sustainability and this behavioral shift was accompanied by higher overall outcome valuation, reduced aversiveness, and a selective level shift for the discounting function for outcomes (but not for aversiveness). These effects were absent under sham stimulation and an active control condition. Together, the findings indicate that sustainable behavior is constrained by a temporally structured value-aversiveness conflict yet can be shifted via non-invasive brain stimulation.

## 1. Introduction

Ensuring environmental sustainability is one of the defining challenges of the twenty-first century^1^. Prior research has emphasized economic and technological determinants of sustainability, whereas the cognitive processes that support or constrain sustainable behaviors remain comparatively underexplored^2^. A prominent example is the knowledge–action gap: despite broad public awareness of environmental risks, a persistent gap remains between this awareness and the consistent adoption of sustainable behaviors and lifestyles needed to mitigate these risks^3–5^. We propose that this gap reflects not primarily deficits in knowledge or social norms, but fundamental cognitive and computational mechanisms that govern sustainable decision-making, particularly temporal discounting^6,7^.

Conventional interventions to promote sustainable behaviors often emphasize long-term environmental benefits (e.g., “*We do not inherit the Earth from our ancestors; we borrow it from our children*” and “*Go green for future generations*”). Yet such strategies may fail because delayed benefits are systematically devalued, a hallmark of temporal discounting^6,7^. Moreover, emphasizing benefits alone is insufficient because decisions require weighing benefits against accompanying costs^8,9^. In sustainability contexts, immediate inconvenience and effort costs frequently outweigh delayed benefits (e.g., *waste sorting requires learning sorting rules and adding extra steps at the point of disposal*); consequently, highlighting long-term benefits without addressing immediate costs may inadvertently reinforce preferences for unsustainable behaviors.

Existing accounts of sustainable behaviors generally fall into several frameworks. Attitudinal and normative models emphasize cognitive evaluations and social influences; for example, the Theory of Planned Behavior (TPB) links intentions to attitudes, norms, and perceived behavioral control^10,11^, but it often leaves substantial variance in behaviors unexplained and struggles to account for the intention–behaviors gap when external barriers are present^12,13^. Value-based approaches, such as Value–Belief–Norm (VBN) theory, link values to norms through beliefs about consequences and responsibility, and can predict altruistically motivated actions (e.g., conservation) but are less effective for self-interested or habitual behaviours^14^. Motivation-focused models, including Self-Determination Theory (SDT), distinguish intrinsic from extrinsic motivations and suggest that autonomous motivation supports sustained pro-environmental action, whereas extrinsic incentives can undermine long-term commitment^15,16^. Threat-based frameworks such as Protection Motivation Theory (PMT) propose that protective behaviors depend on threat appraisal and coping efficacy, particularly when perceived costs are low^17,18^. However, these frameworks often do not fully explain why sustainable intentions fail to translate into action when decisions involve immediate costs and temporally delayed values.

To address this gap, we propose and test the Temporal-Value Decision Model (TVDM), a quantitative framework for modelling the decision processes that underlie sustainable behaviors. The TVDM separates three components: (i) the subjective value of delayed environmental outcomes, modelled via a discounting function; (ii) the immediate aversiveness (inconvenience/effort cost) of sustainable actions; and (iii) the dynamic interaction between these processes that ultimately guides choice. Temporal discounting describes how individuals systematically devalue rewards that are delayed in time^6,7^, and has been used to explain behaviors in which future benefits are sacrificed for immediate gratification (e.g., unhealthy eating^19,20^, addiction^21^, and procrastination^22,23^). Because the benefits of sustainability (e.g., a stable climate, clean air, and healthy ecosystems) often accrue decades or generations later, their subjective value is strongly discounted. Consistent with this, work on intergenerational discounting in sustainability decisions shows that discounting reduces climate-related cooperation and relates to identifiable neural proceseses^24–26^. Critically, temporal discounting alone does not fully explain unsustainable behaviors. The TVDM therefore incorporates a second barrier: the immediate aversiveness of sustainable actions. Many sustainable behaviors require extra effort or inconvenience (e.g., biking instead of driving, recycling, or using public transport), introducing immediate physical or psychological costs that favor unsustainable alternatives. Finally, the TVDM formalizes the central obstacle to sustainable behaviors as the interaction between immediate aversiveness and delayed, discounted value: when discounted future value outweighs immediate cost, sustainable actions are more likely; otherwise, behaviors are constrained. By jointly modelling value evaluation, delay discounting, and aversiveness, the TVDM targets mechanisms that are not simultaneously captured in prior frameworks.

While the TVDM provides a principled account of the temporal–value trade-off in sustainability decisions, its neural computations and the causal role of specific brain regions remain underspecified^27,28^. Sustainable decision-making likely recruits interacting neural networks, making it essential to identify key nodes that implement TVDM computations and could serve as targets for intervention. Converging evidence implicates prefrontal regions—notably dorsolateral (DLPFC) and ventromedial (vmPFC) subregions—in valuation and self-control processes relevant to sustainability^29,30^. and computational work on temporal discounting highlights contributions of these prefrontal systems^31–35^. Specifically, baseline lateral prefrontal cortex activation (measured by EEG)^36^ and cortical thickness in the vmPFC and DLPFC^37^ have been linked to everyday pro-environmental choices and intergenerational preferences. Neuroimaging paradigms integrating memory control^38,39^ with sustainable decision-making further suggest dissociable roles: with vmPFC activity supporting intentions for sustainable behaviors (potentially via perspective thinking) and DLPFC activity associated with inhibiting unsustainable actions, consistent with its role in cognitive control^40^. However, causal evidence from non-invasive stimulation is mixed: cathodal high-definition transcranial direct current stimulation (HD-tDCS) over left DLPFC has been reported to increase environmentally sustainable decision-making ^41^, whereas inhibitory theta-burst stimulation of right DLPFC has shown no detectable effect on sustainable decision-making^42^. Stimulation of the right temporoparietal junction (rTPJ), a region implicated in perspective taking, has also been reported to increase sustainable decision-making^43^. To adjudicate between these accounts, we directly modulated DLPFC activity using HD-tDCS and paired stimulation with temporal-value decision model (TVDM)-based computational analyses. As a non-invasive and comparatively low-cost technique, tDCS is particularly attractive as a complement to other brain-stimulation interventions. This approach not only tests whether DLPFC modulation shifts sustainable decisions but also identifies the computational route by which it does so. Specifically, whether stimulation reduces sensitivity to immediate task aversiveness, increases the subjective valuation of delayed outcomes, and/or alters temporal discounting. Despite heterogeneity in the literature, we targeted the left DLPFC for three reasons. First, sustainable and prosocial decisions plausibly rely on overlapping neural computations of control and valuation^44^, and meta-analytic evidence suggests that tDCS over DLPFC can increase prosocial behavior^45^. Second, DLPFC stimulation has been shown to influence temporal discounting^46,47^, and reduced procrastination^48,49^, processes conceptually aligned with the TVDM. Third, as a core hub of executive control networks, the DLPFC is a tractable target for potential real-world interventions compared with deeper regions such as vmPFC and offers a means to modulate communication within and between mentalizing and cognitive control networks.

In summary, we combine computational modelling and non-invasive brain stimulation to investigate the neurocomputational mechanisms governing sustainable decision-making. First, we develop and validate the TVDM, which formalizes the trade-off among task aversiveness, outcome value, and temporal discounting in sustainable choice. Second, we test the causal role of DLPFC by applying excitatory tDCS to the left DLPFC; we hypothesize that stimulation will increase sustainable decisions. Critically, we predict that stimulation-induced behavioral shifts will be mediated by selective changes in TVDM parameters (e.g., increased valuation of delayed outcomes, reduced perceived aversiveness, and/or altered discounting). We evaluate above predictions in a behavioral study (Study 1; N = 70) and a two-session brain stimulation study (Study 2; N =128) using the same task and stimulus set. By revealing neurocomputational mechanisms underlying sustainable decision making, our results offer a principled theoretical and empirical basis for interventions that accelerate behavior change toward global sustainability

## 2. Results

### 2.1 Temporal-value decision model of human sustainable behaviors

We proposed the Temporal-Value Decision Model (TVDM) to explain why people—despite awareness of the benefits of sustainable behaviors—often prefer unsustainable behaviors. The model begins by defining a conceptual value space with current value and future value. Sustainable behaviors (e.g., recycling) typically occupy regions of high future value but low, and sometimes negative, current value because they entail immediate inconvenience; by contrast, unsustainable behaviors (e.g., littering) lie in regions of high current value but low, and sometimes negative, future value owing to their present convenience (**Figure 1A**). TVDM embeds this value space within a temporal framework to predict when and why choices shift from unsustainable to sustainable (**Figure 1B**). The model rests on two core cognitive–computational mechanisms. First, an outcome–aversiveness comparison: at decision time T, choice depends on the perceived aversiveness of performing an action at T and the perceived outcome it will yield; a high expected future outcome motivates action only when it outweighs present aversiveness (i.e., Value _outcome_ > Value _aversiveness_). Second, temporal discounting of outcome value: outcomes of sustainable behaviors often happen in the future (T + n) and are therefore discounted at time T (i.e., Value _T_<Value _T+n_), whereas aversiveness is experienced in the present and is relatively unaffected by delay. Consequently, at decision time T, the discounted value of a delayed environmental outcome is often smaller than the immediate aversiveness associated with performing the sustainable action, thereby limiting uptake. Unsustainable actions, which typically have lower present aversiveness, remain preferred despite their low future value. Building on this standard TVDM, we specify four modulatory variants for intervention: to increase sustainable behavior, the Outcome-Enhanced Model (**Figure 1C**) raises discounted future outcome value and the Aversiveness-Inhibited Model (**Figure 1D**) reduces present aversiveness; to decrease unsustainable behavior, the Outcome-Inhibited Model (**Figure 1E**) diminishes future outcome valuation and the Aversiveness-Enhanced Model (**Figure 1F**) increases present aversiveness (for example, by increasing the salience of negative consequences or imposing penalties).

**Figure 1.**
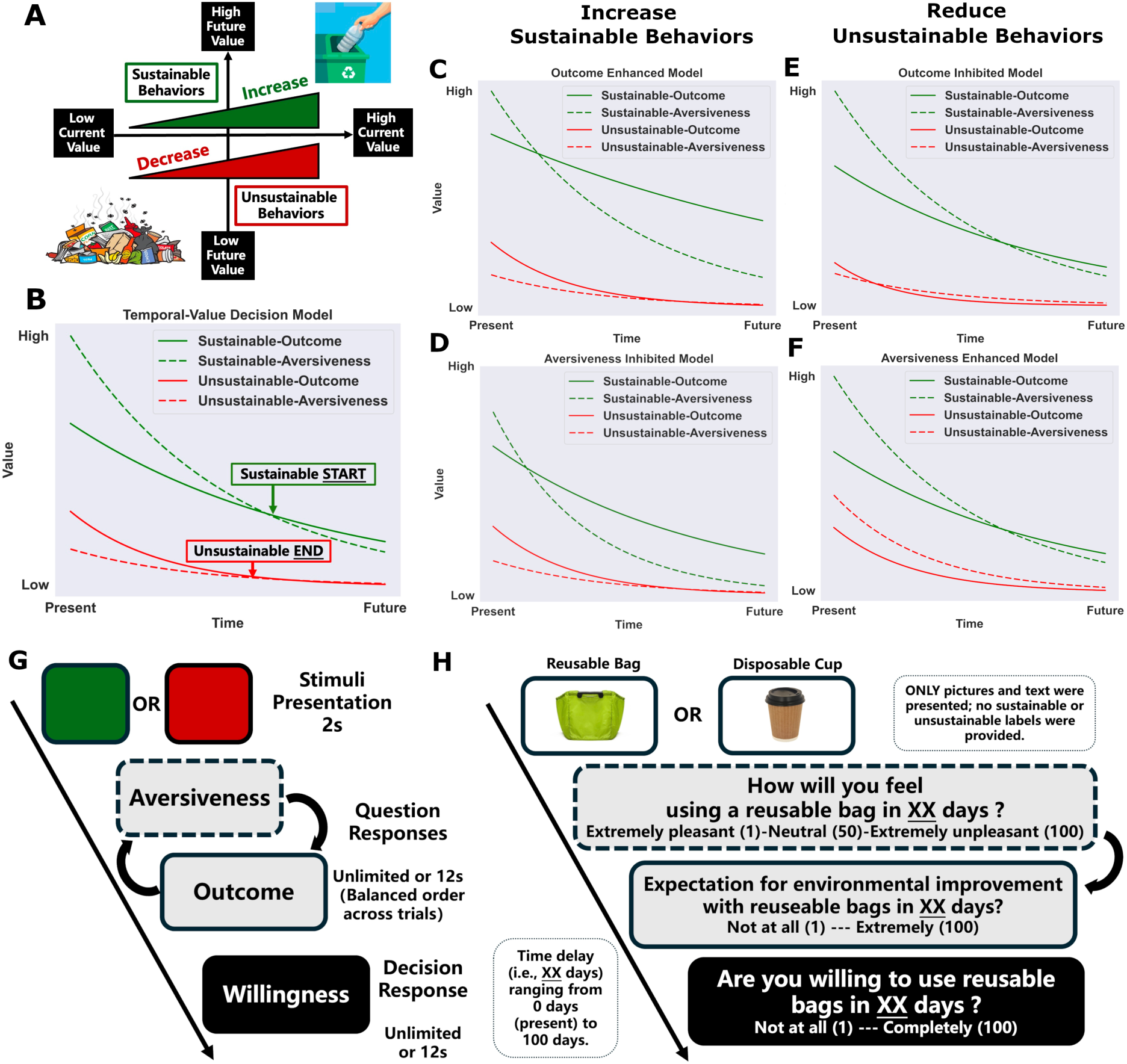
Temporal-Value Decision Model and Task Design. **(A)** Conceptual value space for sustainable and unsustainable behaviors, defined by current value (x-axis) and future value (y-axis). Sustainable behaviors tend to have low (often negative) current value but high future value, whereas unsustainable behaviors show the opposite pattern. **(B)** Standard Temporal-Value Decision Model (TVDM). Temporal functions depict perceived outcome value and aversiveness for sustainable (green) and unsustainable (red) behaviors; “Sustainable START” and “Unsustainable END” mark time points at which the balance between discounted outcomes and immediate aversiveness shifts. **(C–D)** Model variants that increase sustainable behavior: the Outcome-Enhanced Model increases discounted outcome value, and the Aversiveness-Inhibited Model reduces present aversiveness. **(E–F)** Model variants that reduce unsustainable behavior: the Outcome-Inhibited Model decreases outcome valuation, and the Aversiveness-Enhanced Model increases present aversiveness. **(G)** Trial structure. Participants viewed a behavior stimulus and then rated aversiveness and outcome, followed by willingness to perform the behavior. **(H)** Example trial. Images and brief text described the behavior without explicit sustainability labels; ratings were provided on 1–100 scales, and the queried time delay varied from 0 (i.e., present) to 100 days later.

To test TVDM, we designed a sustainable decision-making task that measured participants’ perceived aversiveness, perceived outcome, and willingness to act across temporal delays (**Figure 1G**). On each trial, participants viewed an image and brief text describing a certain behavior; critically, no explicit “sustainable/unsustainable” labels were provided. Participants then provided aversiveness/outcome ratings and a willingness rating in sequence, each on a 1–100 scale. Temporal delay was manipulated by querying the same ratings at five delay values per participant, drawn from one of four fixed schedules spanning from the present (0 days) to 100 days in the future (**Figure 1H**).

### 2.2 Validation of stimuli and paradigm for studying sustainable decision making

Before testing the temporal-value decision model (TVDM) on participants’ experimental data, we validated the stimuli and decision-making paradigm. In an independent online study (N = 480) matched to the main laboratory samples on age, sex, education, and socioeconomic background, participants evaluated candidate images depicting sustainable and unsustainable behaviors (*Supplemental Methods and Results*; **Table S1**). All candidate stimuli were drawn from a previous study of the brain networks underlying sustainable behaviours^40^. From this pool, we selected 30 of 72 images for the subsequent decision-making tasks based on higher mean clarity ratings (*F*_mean_(1, 68) = 4.26, *p*_mean_= 0.04, ɳ^2^ = 0.05) and lower standard deviations of clarify ratings across participants (*F*_std_(1, 68) = 4.36, *p*_std_= 0.04, ɳ^2^ = 0.05) relative to discarded images. Critically, without providing sustainability labels, sustainable stimuli were rated as more environmentally friendly than unsustainable stimuli (t(70) =11.90, *p* < 0.001, Cohen’s d=2.80). Selected images also exhibited larger differences in sustainable valence than discarded images (*F*_mean_(1, 68) = 26.96, *p*_mean_< 0.001, *ɳ^2^*_mean_ = 0.094; *F*_std_(1, 68) = 26.47, *p*_std_< 0.001, *ɳ^2^*_std_ = 0.103), supporting their suitability for probing sustainability-related decisions. These stimuli validation steps increase confidence that subsequent results reflect reliable valuation of the stimuli rather than stimulus ambiguity.

Furthermore, task-internal validity was supported by a negative association between willingness for sustainable versus unsustainable behaviors. Individuals who were more willing to engage in sustainable actions were concurrently less inclined toward unsustainable ones (*r*=-0.40, *p*< 0.001, 95% CI= [-0.54, -0.25]). We assessed external (ecological) validity by relating task choices to trait-level pro-environmental tendency measured with the General Ecological Behavior (GEB) questionnaire^50–52^: higher proportion of sustainable decisions correlated positively with GEB scores (*r*_sus_=0.35, *p*_sus_<0.001, 95% CI_sus_= [0.19, 0.50]). whereas a higher proportion of unsustainable choices correlated negatively with GEB scores (*r*_non-sus_=-0.18, *p*_non-sus_=0.04, 95% CI_non-sus_= [-0.34, -0.01] ). Together, these results indicate that the stimuli and paradigm reliably capture individual differences in sustainable decision-making, generalize to trait-level measures of pro-environmental behavior, and provide a sound empirical basis for fitting and interpreting the TVDM.

### 2.3 Temporal delay reweights perceived aversiveness and value of sustainable decisions

We first tested whether perceived outcome value, task aversiveness, and willingness to act differed between sustainable and unsustainable behaviors in Study 1. Sustainable behaviors were rated as having higher outcome values than unsustainable behaviors (t(69) = 18.34, *p* < 0.0001, Cohen’s d = 2.19, 95% CI [31.56, 39.27], **Figure S1A**). By contrast, perceived aversiveness did not differ between behavior types (t(69) = −0.83, *p* = 0.4108, Cohen’s d = −0.10, 95% CI [−6.04, 2.50], **Figure S1A**), and participants reported comparable willingness to perform sustainable versus unsustainable behaviors (t(69) = 0.23, *p* = 0.8173, Cohen’s d = 0.03, 95% CI [−3.16, 4.00], **Figure S1A**). This dissociation suggests that increasing perceived outcomes alone (for example, via public messaging) may be insufficient to increase willingness to act sustainably. In Study 2, we replicated the outcome-value advantage for sustainable behaviors (t(127) = 41.18, *p* < 0.0001, Cohen’s d = 3.64, 95% CI [39.84, 43.86], **Figure S1B**).

In addition, sustainable behaviors were rated as less aversive than unsustainable behaviors (t(127) = −8.48, *p* < 0.0001, Cohen’s d = −0.75, 95% CI [−23.14, −14.39], **Figure S1B**), and participants expressed greater willingness to perform sustainable behaviors (t(127) = 8.79, *p* < 0.0001, Cohen’s d = 0.78, 95% CI [10.58, 16.74], **Figure S1B**). Together, Studies 1–2 indicate that willingness reflects the combined contribution of outcome value and perceived aversiveness: higher outcome values appear most likely to translate into greater willingness when sustainable actions are also perceived as less aversive.

Finally, to evaluate temporal influences, we tested whether increasing delay systematically altered perceived outcome value, perceived aversiveness, and willingness to act, separately for sustainable and unsustainable behaviors. We quantified delay effects as within-participant linear slopes across the five queried delays (time coded on days). For sustainable behaviors in Study 1, greater delay was associated with a reliable decrease in perceived aversiveness (mean slope = −0.116, t(69) = −4.63, *p* < 0.0001, Cohen’s dz = −0.55, 95% CI [−0.167, −0.066]) and a reliable increase in willingness (mean slope = 0.131, t(69) = 4.85, *p* < 0.0001, Cohen’s dz = 0.58, 95% CI [0.077, 0.185]), whereas outcome value showed a trend-level change with delay (mean slope = −0.029, t(69) = −1.81, *p* = 0.074, Cohen’s dz = −0.22, 95% CI [−0.060, 0.003]). In Study 2, for sustainable behaviors, perceived outcome value decreased robustly with delay (mean slope = −0.105, t(127) = −8.78, p < 0.0001, Cohen’s dz = −0.78, 95% CI [−0.129, −0.082]), whereas aversiveness (mean slope = 0.009, t(127) = 0.57, p = 0.5708, Cohen’s dz = 0.05, 95% CI [−0.022, 0.039]) and willingness (mean slope = 0.008, t(127) = 0.54, p = 0.5885, Cohen’s dz = 0.05, 95% CI [−0.022, 0.038]) did not change reliably with delay. Thus, the delay-related increase in sustainable willingness observed in Study 1 was attenuated under the Study 2 timing constraints, motivating the subsequent computational tests that focus on outcome–aversiveness integration.

### 2.4 Study1 Computational evidence: outcome value and aversiveness jointly predict sustainable choice

We used trial-by-trial data to evaluate computational models predicting participants’ willingness to perform sustainable and unsustainable behaviors from perceived outcome value and task aversiveness. We compared four candidate models (**Figure 2A**): Model 1 assumed willingness was driven by outcome value alone, Model 2 by task aversiveness alone, Model 3 combined outcome value and task aversiveness additively, and Model 4 added their interaction. Models were compared using leave-one-out cross-validation (LOO-CV; changes in expected log predictive density, ΔELPD), information criteria (AIC/BIC), and participant-wise cross-validated predictive log-likelihood (see *Supplementary Methods and Results for details*).

**Figure 2.**
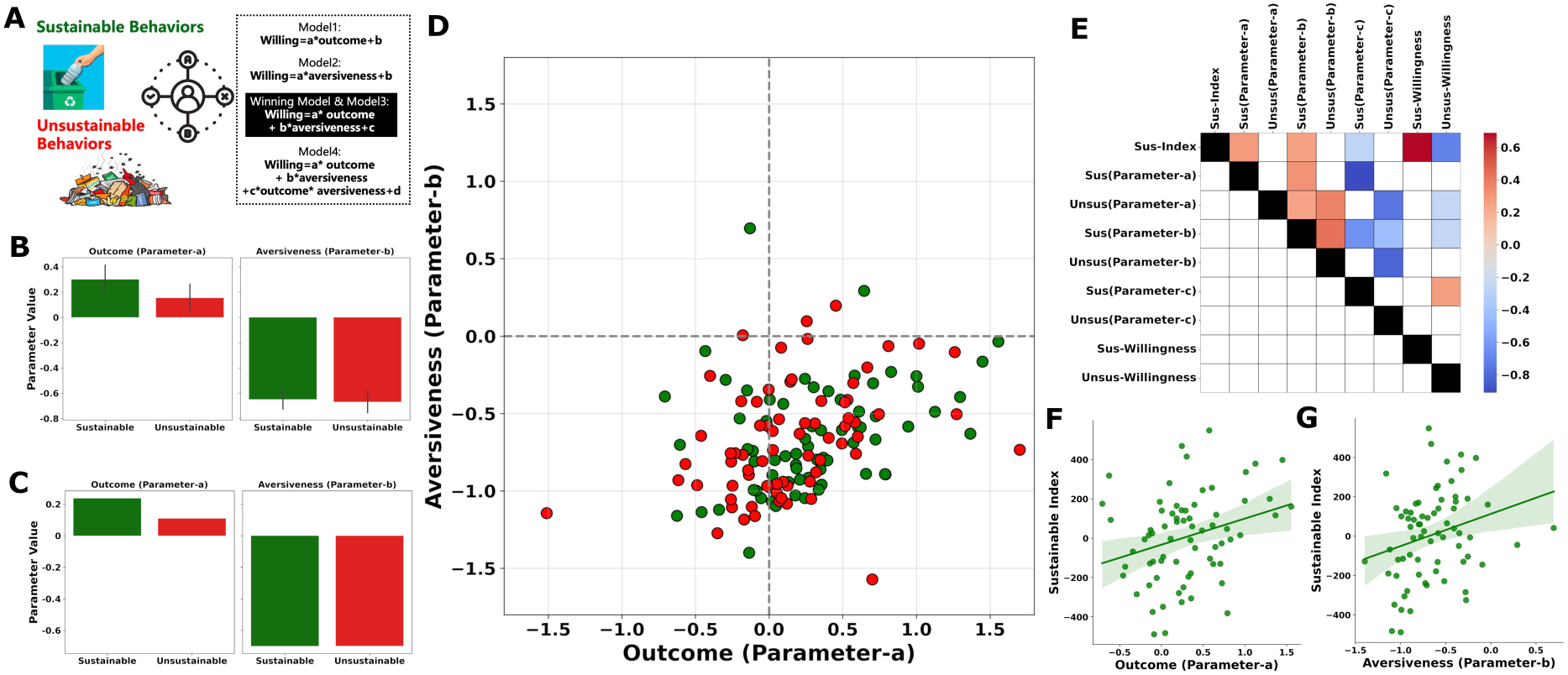
Predictive power of computational models: estimating sensitivity to outcomes and aversiveness for sustainability prediction. **(A)** Four candidate computational models were evaluated to explain decision-making processes underlying sustainable and unsustainable behaviors. Model 3, incorporating outcome and aversiveness factors without interactions, emerged as the best-fitting model (see Supplemental Materials for detailed model comparisons). **(B)** Individual-level comparisons of parameter-a (sensitivity to outcomes) and parameter-b (sensitivity to aversiveness) between sustainable and unsustainable models. **(C)** Group-level comparisons of parameters-a and b between sustainable and unsustainable models. **(D)** Visualization of parameters a and b in a two-dimensional space. Green represents the sustainable model, and red represents the unsustainable model. Positive values of parameter-a indicate a stronger facilitatory effect of outcomes on behavior, while negative values of parameter-b indicate a stronger inhibitory effect of aversiveness on behavior. **(E)** Correlation matrix of all model parameters with willingness to engage in sustainable and unsustainable behaviors. **(F)** A higher value of parameter-a (outcome sensitivity) in the sustainable model is associated with increased overall sustainability. **(G)** A higher value of parameter-b (aversiveness sensitivity) in the sustainable model is also linked to increased overall sustainability.

On sustainable trials, the additive outcome-plus-aversiveness model (Model 3) was preferred (**Figure S2**), outperforming the outcome-only model (ΔELPD = 323.06, SE = 35.39) and the aversiveness-only model (ΔELPD = 28.77, SE = 8.84) and yielding the lowest AIC (9152.78). In Model 3, outcome value positively predicted willingness (β = 0.237, *p* < .001), whereas task aversiveness negatively predicted willingness (β = −0.699, *p* < .001), and both predictors explained unique variance (likelihood-ratio tests: χ² = 23.58 for outcome and 295.28 for aversiveness). Adding an interaction term did not improve predictions for sustainable trials (Model 4; interaction *p* = .656). For unsustainable trials, Model 3 again outperformed Models 1–2 (ΔELPD = 313.70, SE = 32.00; ΔELPD = 5.82, SE = 4.26), and both predictors remained significant (outcome *p* = .022; aversiveness *p* < .001). Although Model 4 yielded a better AIC and a significant interaction (β = 0.0034, *p* = .015), the incremental predictive benefit of the interaction was small. Accordingly, given Model 3’s robust performance across both trial types and evidence that outcome value and aversiveness contribute unique explanatory power, we treat Model 3 as the primary account of willingness.

For interpretability, we summarized Model 3 in terms of (a) an outcome weight (sensitivity to outcome value), (b) an aversiveness weight (sensitivity to task aversiveness), and (c) a decision/bias parameter (e.g., intercept or inverse temperature). We fit Model 3 separately to each participant’s sustainable and unsustainable trials to obtain individual-level parameter estimates. The outcome weight (a) tended to be larger for sustainable than unsustainable trials, although this difference did not reach statistical significance (mean difference = 0.15, 95% CI [−0.02, 0.31], Cohen’s d = 0.21, *p* = .078; **Figure 2B**). No group-level difference was observed for the aversiveness weight (b) (mean difference = 0.02, 95% CI [−0.07, 0.11], Cohen’s d = 0.05, *p* = .666; **Figure 2B**). Despite modest between-participant variability, fitting Model 3 to all participants’ trial-level data reproduced the same qualitative pattern of outcome–aversiveness integration (**Figure 2C**).

To visualize heterogeneity in outcome and aversiveness weights, we plotted each participant’s parameter pair (a, b), estimated separately for sustainable and unsustainable trials, in a two-dimensional parameter space (**Figure 2D**). Most estimates fell in the fourth quadrant (positive outcome weight and negative aversiveness weight), consistent with greater sensitivity to outcome value and reduced sensitivity to aversiveness being associated with higher willingness to choose sustainable options. To assess the psychological validity of these parameters, we related them to a subject-level sustainability index (mean willingness for sustainable items minus mean willingness for unsustainable items). In the correlation matrix relating model parameters to individual measures (**Figure 2E**). Across participants, higher outcome sensitivity (parameter a) was associated with a higher sustainability index (r(69) = 0.29, *p* < .001; **Figure 2F**). Lower aversiveness sensitivity (i.e., parameter b closer to zero / higher b values) was also associated with the index (r(69) = 0.26, *p* < .001; **Figure 2G**). The decision/bias parameter (c) estimated on sustainable trials was negatively associated with the sustainability index (r(68) = −0.27, *p* < .001; **Figure S3**). Notably, parameters estimated from unsustainable trials did not show corresponding associations (*p*s > 0.05). Together, these results indicate that individual differences in the joint weighting of outcome value and task aversiveness (rather than outcome value alone) predict individual differences in sustainable decision-making.

### 2.5 Study1 Asymmetric outcome–aversiveness dynamics differentiate sustainable from unsustainable decisions

To examine how participants integrate perceived outcome and task aversiveness during sustainable decision making, and to test the hypothesis that sustainable actions are most likely when perceived outcomes outweigh aversiveness, we stratified trials into four decision categories (“not to do,” “reluctant,” “consider,” “to do”). Within each category, we quantified (i) the absolute ratings of outcome and aversiveness and (ii) their relative ordering (outcome > aversiveness vs. outcome < aversiveness). We found that task aversiveness exerted a stronger influence on sustainable decisions than outcome value. To express outcome value and aversiveness on a common scale, we estimated the optimal scaling weight for aversiveness relative to outcome value and found that a weight of 1.44 maximized model performance (*see Supplementary Methods and Results*; **Figure S4**). Accordingly, we scaled aversiveness ratings by 1.44 in subsequent analyses to directly compare the effects of aversiveness and outcome value on choice.

Sustainable trials conformed to this proposed integration account. A repeated-measures ANOVA showed that the outcome–aversiveness relationship depended strongly on decision category, yielding a robust interaction (F(3, 162) = 101.77, *p* < .001, ηp² = .653, **Figure 3A**). In the low-willingness categories (“not to do” and “reluctant”), perceived aversiveness exceeded outcome, consistent with declining sustainable actions when perceived costs outweighed benefits (not to do: M_Outcome_ = 58.17 vs. M_Aversiveness_ = 101.71, M_diff_ = −43.54, t(57) = −8.34, *p* < .001, Cohen’s d = 1.09; reluctant: M_Outcome_ = 62.01 vs. M_Aversiveness_ = 88.82, M_diff_ = −26.82, t(68) = −11.20, *p* < .001, d = 1.35). As willingness increased (“consider” and “to do”), perceived aversiveness decreased and perceived outcome increased, producing a crossover such that outcome exceeded aversiveness in the high-willingness categories (consider: M_Outcome_ = 67.19 vs. M_Aversiveness_ = 61.28, M_diff_ = 5.91, t(68) = 2.51, *p* = .015, Cohen’s d = 0.30; to do: M_Outcome_ = 75.79 vs. M_Aversiveness_ = 36.88, M_diff_ = 38.91, t(66) = 11.80, *p* < .001, Cohen’s d = 1.44).

**Figure 3.**
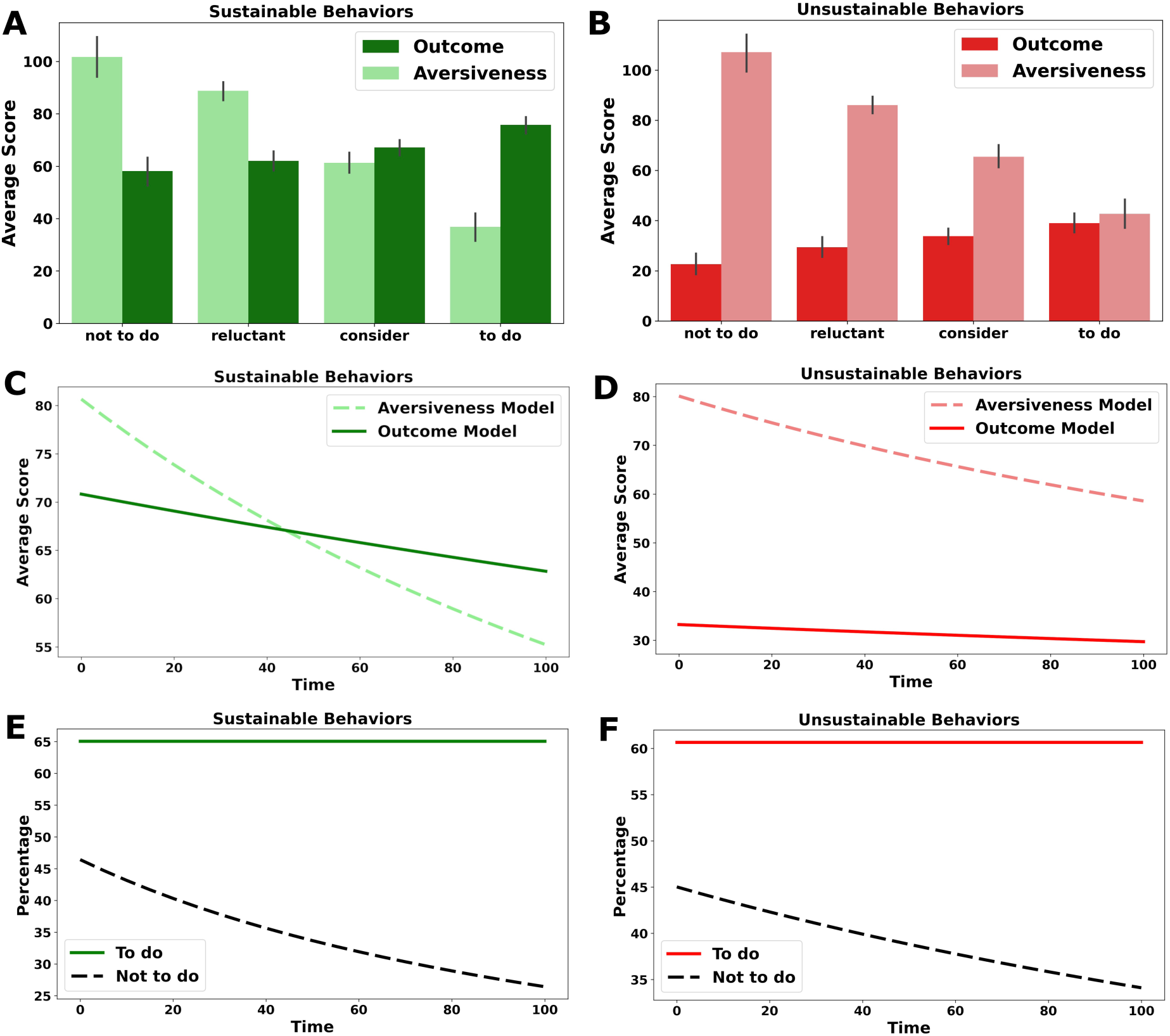
Temporal dynamics of outcome-aversiveness comparisons led to different sustainable decisions. **(A)** Individuals choose against sustainable behaviors when task aversiveness exerts a stronger influence than outcomes, but they choose to engage in behaviors when aversiveness is less influential than outcomes. **(B)** Outcome-aversiveness comparisons do not affect decisions regarding unsustainable behaviors, despite a significant decrease in aversiveness from the option not-to-do to option to-do. **(C)** Sustainable behaviors: fitted trajectories of task aversiveness (dashed) and outcome (solid) across time. Aversiveness declines more steeply than outcome, producing a crossover in which outcome exceeds aversiveness at later time points. **(D)** Unsustainable behaviors: fitted trajectories of aversiveness (dashed) and outcome (solid) across time. Although both signals decrease over time, outcome remains lower than aversiveness across the time horizon. **(E)** Sustainable behaviors: model-derived distribution of responses classified as “to do” (solid) versus “not to do” (dashed) across time. Although the proportion of individuals choosing sustainable behaviors does not vary over time, the decision not to engage in sustainable actions is highest in the present and decreases as time delay increases. (F) Unsustainable behaviors: analogous model-derived response distribution across time. The percentage of individuals choosing unsustainable behaviors remains consistent over time, but the decision not to engage in such behaviors is highest in the present and decreases as time delay increases.

By contrast, unsustainable trials showed the opposite integration profile: outcome remained lower than aversiveness across decision categories, even when participants reported greater willingness to engage in unsustainable behaviors. A repeated-measures ANOVA confirmed that the outcome–aversiveness relationship varied by decision category (F(3, 123) = 61.16, *p* < .001, ηp² = .599; **Figure 3B**). Descriptively, outcome increased with willingness (not to do: M_Outcome_ = 22.62; reluctant: M_Outcome_ = 29.41; consider: M_Outcome_ = 33.83; M_Outcome_ = to do: 39.03), whereas aversiveness decreased (not to do: M_Aversiveness_ = 107.21; M_Aversiveness_ =reluctant: 86.10; M_Aversiveness_ =consider: 65.51; M_Aversiveness_ = to do: 42.73). However, planned paired comparisons indicated that aversiveness reliably exceeded outcome in the three lower-willingness categories (not to do: M_diff_ = −84.59, t(56) = −16.93, *p* < .001, Cohen’s d = 2.24; reluctant: M_diff_ = −56.69, t(62) = −20.69, *p* < .001, Cohen’s d = 2.61; consider: M_diff_ = −31.69, t(65) = −11.39, *p* < .001, Cohen’s d = 1.40). Notably, even in the highest-willingness category (“to do”), outcome did not exceed aversiveness; instead, the two ratings converged and the difference was not significant (M_diff_ = −3.70, t(60) = −1.11, *p* = .269, Cohen’s d = 0.14), indicating that increased willingness to engage in unsustainable actions did not reflect a clear subjective advantage of outcomes over aversiveness.

Extending these category-based analyses, we tested whether the relative weighting of perceived outcome and task aversiveness unfolds dynamically over the time delay. For sustainable behaviors, the individual-level discounting rate for aversiveness was significantly greater than that for outcome (t(69) = 3.84, *p* < 0.001; Wilcoxon signed-rank W = 547, *p* < 0.001; **Figure 3C**), indicating that aversiveness diminished more rapidly over time. Consequently, the balance between the two signals shifted, producing a crossover in which outcome exceeded temporally weighted aversiveness at later delays (group-level crossover at approximately t ≈ 44 on the 0–100 scale). This crossover pattern was present in a majority of participants (65.7%), consistent with systematic temporal reweighting favoring delayed outcomes in sustainable decisions.

For unsustainable behaviors, the temporal profile differed. Although aversiveness also declined over time and did so more steeply than outcome (t(69) = 2.46, p = 0.017; Wilcoxon W = 636, p < 0.001; **Figure 3D**), perceived outcome remained uniformly low across delays. Accordingly, no group-level crossover between outcome and weighted aversiveness emerged within the observed time horizon, indicating that increased willingness to engage in unsustainable actions cannot be explained by a robust temporal reversal in the relative valuation of outcomes versus aversiveness. Importantly, discounting rates for outcome and aversiveness did not differ significantly between sustainable and unsustainable conditions (all *p*s > 0.41), suggesting that the divergence primarily reflects differences in relative signal magnitudes rather than overall discounting strength.

Consistent with these signal dynamics, model-derived choice distributions showed that the proportion of “not to do” responses declined over time for both sustainable and unsustainable behaviors, whereas “to do” responses remained comparatively stable. Hyperbolic fits to the choice proportions indicated a clear temporal decrease in “not to do” responses in both conditions (sustainable: k = 0.0076, **Figure 3E**; unsustainable: k = 0.0032, **Figure 3F**), while “to do” responses showed negligible discounting (k ≈ 0 in both cases).

Finally, to further characterize how temporal valuation differs across decision outcomes, we refit the hyperbolic trajectories separately for trials culminating in “to do” versus “not to do” decisions. For sustainable behaviors, “to do” trials exhibited high and only weakly discounted outcome values (pooled fit: V_0_ = 82.71, k = 0.00099) alongside lower but more steeply declining aversiveness (V_0_ = 31.24, k = 0.00608), yielding immediate dominance of outcome over aversiveness (unweighted crossover at t ≈ 0; **Figure 4A**). By contrast, sustainable “not to do” trials showed high, essentially non-discounted aversiveness (V_0_ = 74.38, k ≈ 0) paired with more strongly discounted outcomes (V_0_ = 66.98, k = 0.00369), such that outcome never exceeded aversiveness (no crossover; **Figure 4B**). A similar dissociation was observed for unsustainable behaviors. “To do” trials were marked by low, essentially flat outcome (V_0_ = 36.66, k ≈ 0;) and a modestly declining aversiveness signal (V_0_ = 30.35, k = 0.00200), producing a small outcome advantage already at t ≈ 0 **(Figure 4C)**. In “not to do” trials, aversiveness remained high with minimal temporal change (V_0_ = 79.18, k = 0.00059) while outcome stayed uniformly low and flat (V_0_ = 20.32, k ≈ 0; **Figure 4D**).

**Figure 4.**
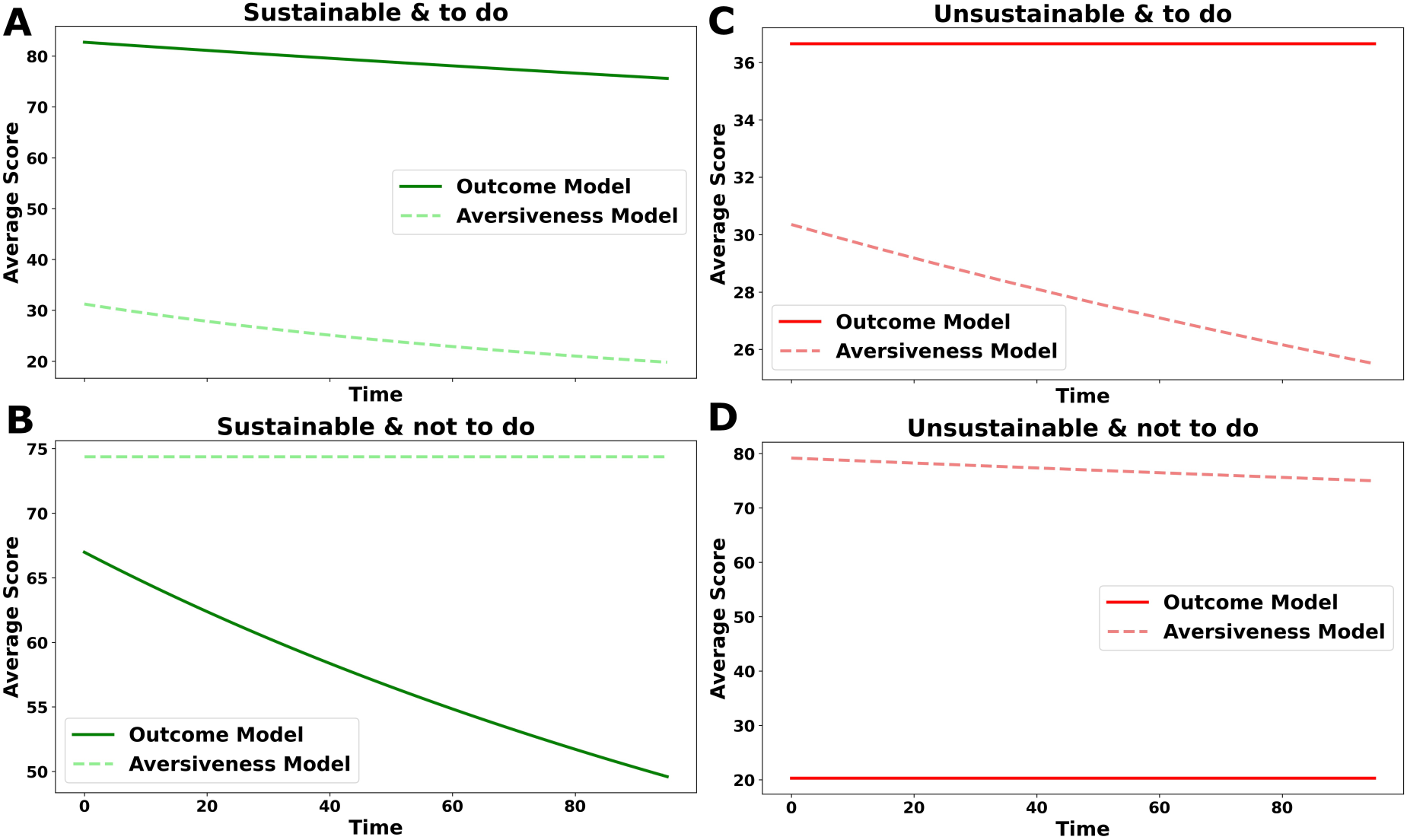
Outcome and aversiveness discounting trajectories dissociate by decision outcome. Hyperbolic discounting functions were fit separately for trials ending in “to do” versus “not to do” choices and plotted for sustainable (green) and unsustainable (red) behaviors. For sustainable behaviors, “to do” decisions were characterized by consistently high outcome values and lower, more steeply discounted aversiveness (A), whereas “not to do” decisions showed persistently high aversiveness with outcomes that declined over time (B). For unsustainable behaviors, “to do” decisions exhibited low, relatively flat outcome values alongside modestly decreasing aversiveness (C), while “not to do” decisions showed sustained high aversiveness with uniformly low outcomes (D).

### 2.6 Study 2 replicates behavioral and model-based predictors of sustainable decisions

Study 1 suggested that: (1) willingness to act sustainably reflects both outcome value and task aversiveness, with a two-factor model (Model 3) outperforming single-factor alternatives; (2) outcome value is weighted more heavily in decisions to act sustainably than in decisions to act unsustainably; (3) greater sensitivity to outcomes and lower sensitivity to aversiveness predict higher individual-level sustainability; and (4) sustainable choices arise primarily when perceived outcomes exceed perceived aversiveness. We tested the robustness of these results in the baseline session of Study 2 (N = 128), which used the same task under a 12-s response limit. We treated this baseline session as an independent, non-stimulation replication sample and tested whether modest time pressure altered the behavioral patterns observed in Study 1.

First, consistent with Study 1, Model 3 again provided the best balance of predictive accuracy and parsimony for both sustainable and unsustainable decisions (*see Supplementary Methods and Results*; **Figure S5**). Second, group-level comparisons of Model 3 parameters showed no reliable differences between sustainable and unsustainable trials in outcome weighting (parameter a: t(127) = 1.30, *p* = 0.19); and parameter (b)) or aversiveness weighting (parameter b: t(127) = 1.01, *p* = 0.31), diverging from Study 1 (**Figure S6**). Thirdly, the individual-difference correlations observed in Study 1 were preserved in Study2 (**Figure S7A**): outcome sensitivity (a) correlated positively with the sustainability index (r(126) = 0.17, *p* = 0.05; **Figure S7B**), reduced aversiveness sensitivity (indexed by higher b) correlated with the index (r(126) = 0.37, *p* < 0.001; **Figure S7C**), whereas parameter (c) correlated negatively with the index (r(126) = −0.21, *p* = 0.02; **Figure S8**). Finally, and critically, category-based outcome–aversiveness ordering patterns in Study2 qualitatively matched Study 1. For sustainable trials, low-willingness responses were characterized by aversiveness exceeding outcomes, whereas higher-willingness responses showed the reverse pattern (**Figures S9**). Accordingly, net value (D) increased monotonically across decision categories and shifted from negative to positive as participants moved from refusal to action. For unsustainable trials, perceived outcomes increased and aversiveness decreased with willingness, but aversiveness remained higher than outcomes across categories, such that D became less negative yet did not cross into positive territory on average. Together, these findings indicate that a 12-s decision-time limit did not change the core behavioral and model-based signatures reported in Study 1, supporting their replicability under modest time pressure in an independent sample (i.e., Study2).

### 2.7 Study2 left DLPFC stimulation shifted decisions toward sustainability

Next, we tested whether excitatory HD-tDCS over the left dorsolateral prefrontal cortex (lDLPFC) causally promotes sustainable decisions. A total of 128 participants completed the same decision-making paradigm as in Study 1 while receiving one of three stimulation conditions (**Figure 5A**): anodal lDLPFC (active-DLPFC; n=43), sham lDLPFC (sham-DLPFC; n=41), or an active control montage over occipital cortex (active-Oz; n=44). tDCS modulates cortical excitability via weak scalp currents; here we applied anodal (putatively excitatory) stimulation to enhance prefrontal activity hypothesized to support future-oriented valuation and inhibitory control. To account for baseline differences in willingness, analyses focused on change scores (Day 3 − Day 1), with Day 1 serving as the pre-stimulation baseline for all participants. To rule out alternative explanations for any stimulation-related effects, we conducted several control analyses. Neither active nor sham stimulation produced reliable changes in self-reported emotion, and groups did not differ in their guesses about stimulation intensity, indicating successful blinding. No severe discomfort events were reported, and self-reported discomfort ratings were comparable across groups (Supplemental Methods and Results; Table S2).

**Figure 5.**
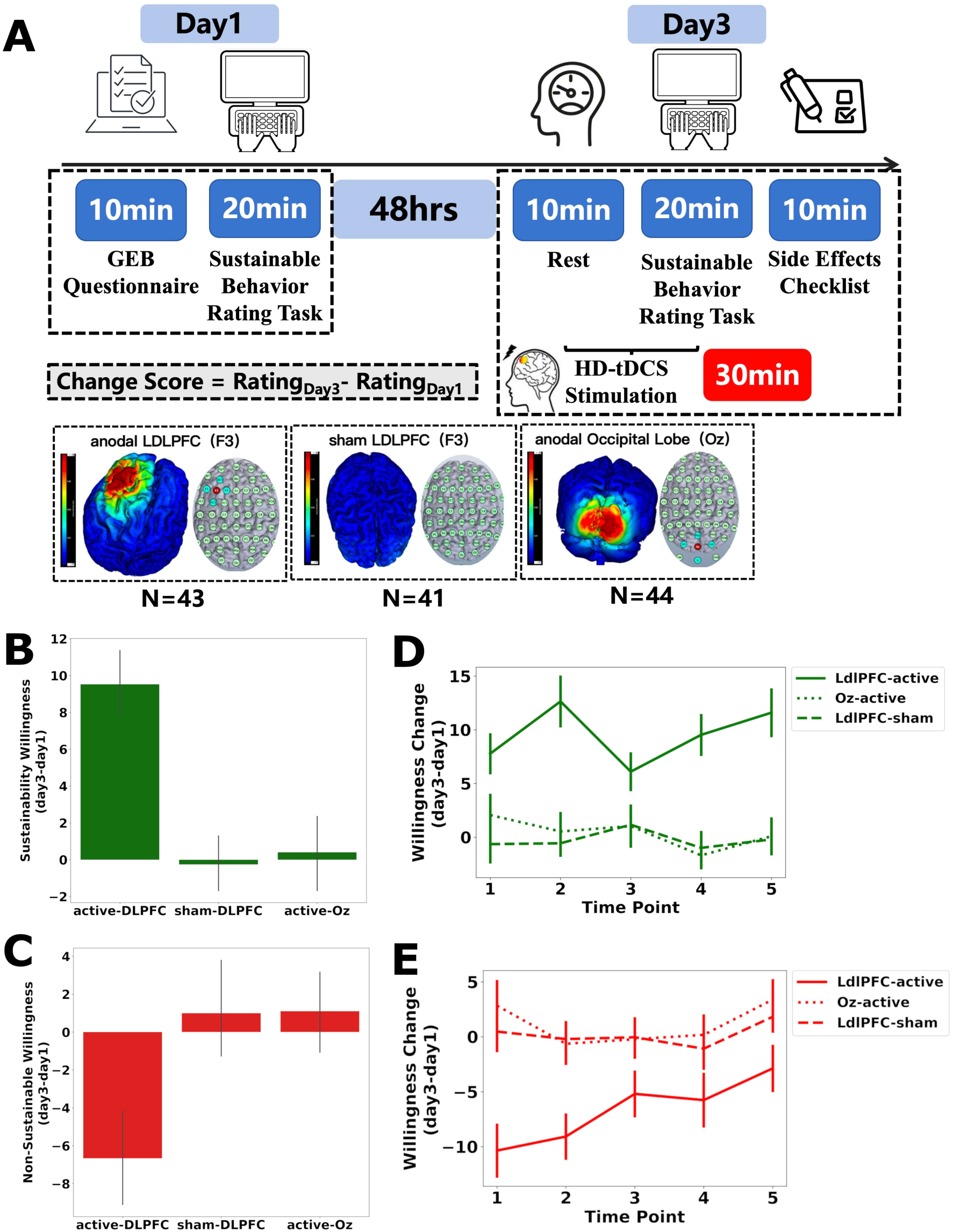
Anodal tDCS over DLPFC enhances sustainable and suppresses unsustainable behaviors. **(A)** Study procedure and stimulation conditions. Electric-field simulations illustrate the three montages: anodal lDLPFC, sham lDLPFC, and anodal occipital cortex. **(B)** Change in willingness to engage in sustainable behaviors across stimulation groups. Relative to sham stimulation and an active control site (Oz), active stimulation over lDLPFC increased willingness to engage in sustainable behaviors. **(C)** Change in willingness to engage in unsustainable behaviors across stimulation groups. **(D)** Sustainable willingness changes as a function of within-trial delay (time points 1–5) for each stimulation group. **(E)** Unsustainable willingness changes as a function of within-trial delay (time points 1–5) for each stimulation group.

Willingness to engage in sustainable behaviors differed robustly across stimulation groups (F(2,125)=35.503, *p*<0.001, η²=0.362; **Figure 5B**). The active-DLPFC group showed a marked increase (mean±s.d., 9.529±6.242), whereas changes were near zero in the active-Oz (0.402±6.592) and sham-DLPFC (−0.255±4.987) groups. Tukey-corrected post hoc tests confirmed that the active-DLPFC group increased more than both active-Oz (mean difference=9.127, 95% CI [6.076, 12.179], *p*<0.001) and sham-DLPFC (mean difference=9.784, 95% CI [6.678, 12.890], *p*<0.001), with no difference between active-Oz and sham-DLPFC (*p*=0.869). A complementary pattern emerged for willingness to engage in unsustainable behaviors (F(2,125)=13.172, *p*<0.001, η²=0.174; **Figure 5C**). Participants receiving active-DLPFC stimulation showed a reduction (−6.660±8.124), whereas the active-Oz (1.103±7.152) and sham-DLPFC (1.000±8.792) groups showed nearly unchanged. Tukey tests indicated a larger decrease under active-DLPFC than under active-Oz (mean difference=−7.764, 95% CI [−11.849, −3.678], *p*<0.001) and sham-DLPFC (mean difference=−7.661, 95% CI [−11.819, −3.502], *p*=0.0001), with no difference between active-Oz and sham-DLPFC (*p*=0.998). Together, these results indicate that anodal stimulation of the lDLPFC—relative to both sham stimulation and an active control site—selectively shifted behavioral willingness toward sustainability.

Because the paradigm manipulates outcome delay within trials, we next tested whether stimulation effects depended on temporal delay (immediate versus delayed decisions). For sustainable trials, a repeated-measures ANOVA including within-trial delay revealed no group × delay interaction (F(8,500)=1.542, *p*=0.140; **Figure 5D**), indicating that the enhancement produced by active-DLPFC stimulation was comparable across delays and evident even at the shortest delay. For unsustainable trials, an analogous analysis likewise yielded no group × delay interaction (F(8,500)=1.137, *p*=0.337; **Figure 5E**), suggesting that the effect of active-DLPFC stimulation was similarly consistent across time delays.

Finally, we conducted exploratory correlational analyses within the active-DLPFC group to probe individual differences in stimulation responsiveness. We correlated baseline willingness (Day 1) with stimulation-induced change (Day 3 − Day 1). Higher baseline willingness to act sustainably was associated with smaller stimulation-related increases in sustainable willingness (r(42)=−0.34, *p*=0.027) and smaller increases in the perceived outcomes of sustainable behaviors (r(42)=−0.33, *p*=0.033). Similarly, higher baseline willingness to act unsustainably was associated with a larger stimulation-related decrease in sustainable willingness (r(42)=−0.32, *p*=0.034). Although exploratory, these associations suggest that stimulation effects were more pronounced among participants showing lower initial sustainability.

### 2.8 Study2 Active stimulation over DLPFC altered underlying TVDM computations

We next examined candidate cognitive–computational mechanisms through which anodal tDCS increased sustainable behavior by comparing key TVDM variables and model parameters across groups, focusing on Day 3 (i.e., stimulation session) − Day 1 (i.e., baseline session) change.

Firstly, we tested whether stimulation altered the evaluative components (i.e., perceived Outcome and Aversiveness) hypothesized to support behavioral change. For sustainable behavior’s, perceived outcome differed by group (F(2,125)=22.496, *p*<0.001, η²=0.265; **Figure S10A**): active-DLPFC increased sustainable outcome (4.895±5.010) relative to active-Oz (0.789±3.366; mean difference=4.105, 95% CI [1.997, 6.213], *p*<0.001) and sham-DLPFC (−1.000±3.891; mean difference=5.895, 95% CI [3.749, 8.040], *p*<0.001). Sustainable aversiveness also differed across groups (F(2,125)=8.154, *p*=0.0005, η²=0.115; **Figure S10B**): active-DLPFC reduced sustainable aversiveness (−4.526±7.448) compared with both active-Oz (0.739±7.714; mean difference=−5.265, 95% CI [−8.810, −1.720], *p*=0.0017) and sham-DLPFC (0.743±5.437; mean difference=−5.269, 95% CI [−8.877, −1.660], *p*=0.0021). Active-Oz and sham-DLPFC did not differ for either sustainable outcome or aversiveness (both *ps*>0.10). For unsustainable behavior’s, active-DLPFC stimulation primarily affected perceived outcome: group differences were significant (F(2,125)=10.141, *p*=0.0001, η²=0.140; **Figure S10C**), with a reduction under active-DLPFC (−4.561±6.816) relative to active-Oz (0.494±4.353; mean difference=−5.055, 95% CI [−7.908, −2.202], *p*=0.0001) and sham-DLPFC (−0.287±5.399; mean difference=−4.275, 95% CI [−7.179, −1.371], *p*=0.0019). By contrast, unsustainable aversiveness was unaffected (F(2,125)=0.291, *p*=0.748, η²=0.005; **Figure S9D**). We additionally evaluated tDCS effects on outcome and aversiveness across time points (See *Supplemental Methods and Results*; **Figure S11**).

Secondly, we fit Model 3, which combines aversiveness and outcome to predict willingness to act, separately for sustainable and unsustainable behaviors. We evaluated tDCS effects on Model3 parameters (See Supplemental Methods and Results for details). Only one effect emerged: in the unsustainable condition, changes in parameter C differed across groups (F(2,125)=5.632, p=.005, η²=0.083, ω²=0.067; means_active-DLPFC_=−10.7±37.9; means_active-Oz_=15.1±33.8; means_sham-DLPFC_=6.1±37.1). Follow-up pairwise comparisons indicated that parameter c decreased under active lDLPFC stimulation compared to active-Oz (mean difference=25.797, 95% CI [10.441, 41.153], *p*=.0013) and sham-DLPFC (mean difference=16.828, 95% CI [0.530, 33.127], *p*=.0432). No group effects were observed for parameters a (outcome sensitivity) or b (aversiveness sensitivity), suggesting that stimulation-related behavioral change did not arise from altered choice sensitivity to outcome or aversiveness per se.

Critically, we asked how changes neural excitability in the DLPFC related to proposed TVDM computational decision mechanisms underlying sustainable behaviors. To characterize how ratings varied with time delay, we first compared a constant model with three time-varying functions (linear, generalized hyperbolic, generalized exponential). For the outcome measure, the generalized exponential model was decisively preferred (AIC = 79.81; BIC = 82.80; R^2^=0.84; ΔAIC = 0; w=0.99; **Figure 6A**). The linear model showed substantially weaker support (AIC = 92.61; BIC = 94.60; 𝑅^2^=0.66; ΔAIC = 12.79; w=0.001658), as did the generalized hyperbolic model (AIC = 93.2840; BIC = 96.2712; 𝑅^2^=0.68; ΔAIC = 13.46; w=0.001187). The constant model performed worst (AIC = 112.4760; BIC = 113.4720; 𝑅^2^=0; ΔAIC = 32.65; 𝑤=8.07×10^−8^). For aversiveness, in contrast, the constant model was best supported (AIC = 117.21; BIC = 118.20; 𝑅^2^=0; ΔAIC = 0; 𝑤=0.58; **Figure 6B**). Using the winning functional forms, we fit each participant’s delay-dependent outcome and aversiveness (separately for sustainable and unsustainable behaviors) and tested whether stimulation altered parameter changes from Day 1 to Day 3. Across sustainable outcome fits, change in the intercept parameter y0 significantly differed between stimulation group predicted (F(2,125)=9.62, *p*<.001, η²=0.133, ω²=0.119; **Figure 6C**). Relative to both control groups, active-DLPFC showed a larger positive shift in y0 (means_active-DLPFC_=4.47; means_active-Oz_=0.30; means_sham-DLPFC_=−1.71), with pairwise differences for active-DLPFC vs active-Oz (t_Welch_=2.89, *p*=.005, d=0.62) and active-DLPFC vs sham-DLPFC (t_Welch_=4.10, *p*=.0001, d=0.89); active-Oz did not differ from sham (*p*=.137). In contrast, no group effects were observed for changes in the exponential rate (k) or offset/asymptote (c) (both *p*>.20), indicating that DLPFC stimulation primarily shifted the overall level of outcome valuation rather than the temporal sensitivity of discounting. Critically, we showed that the tDCS effect was valence and context-dependent. For unsustainable outcomes, stimulation group differed in Δy0 (F(2,125)=5.07, *p*=.008, η²=0.075, ω²=0.060; **Figure 6D**), but in the opposite direction: active-DLPFC showed a more negative shift (means_active-DLPFC_=−5.84; means_active-Oz_=−1.48; means_sham-DLPFC_=−0.60), differing from active-Oz (t_Welch_=−2.49, *p*=.0147, d=0.53) and sham-DLPFC (t_Welch_=−2.87, *p*=.0052, d=0.63). Again, Δk and Δc were not modulated by brain stimulation (*p*’s>.79).

**Figure 6.**
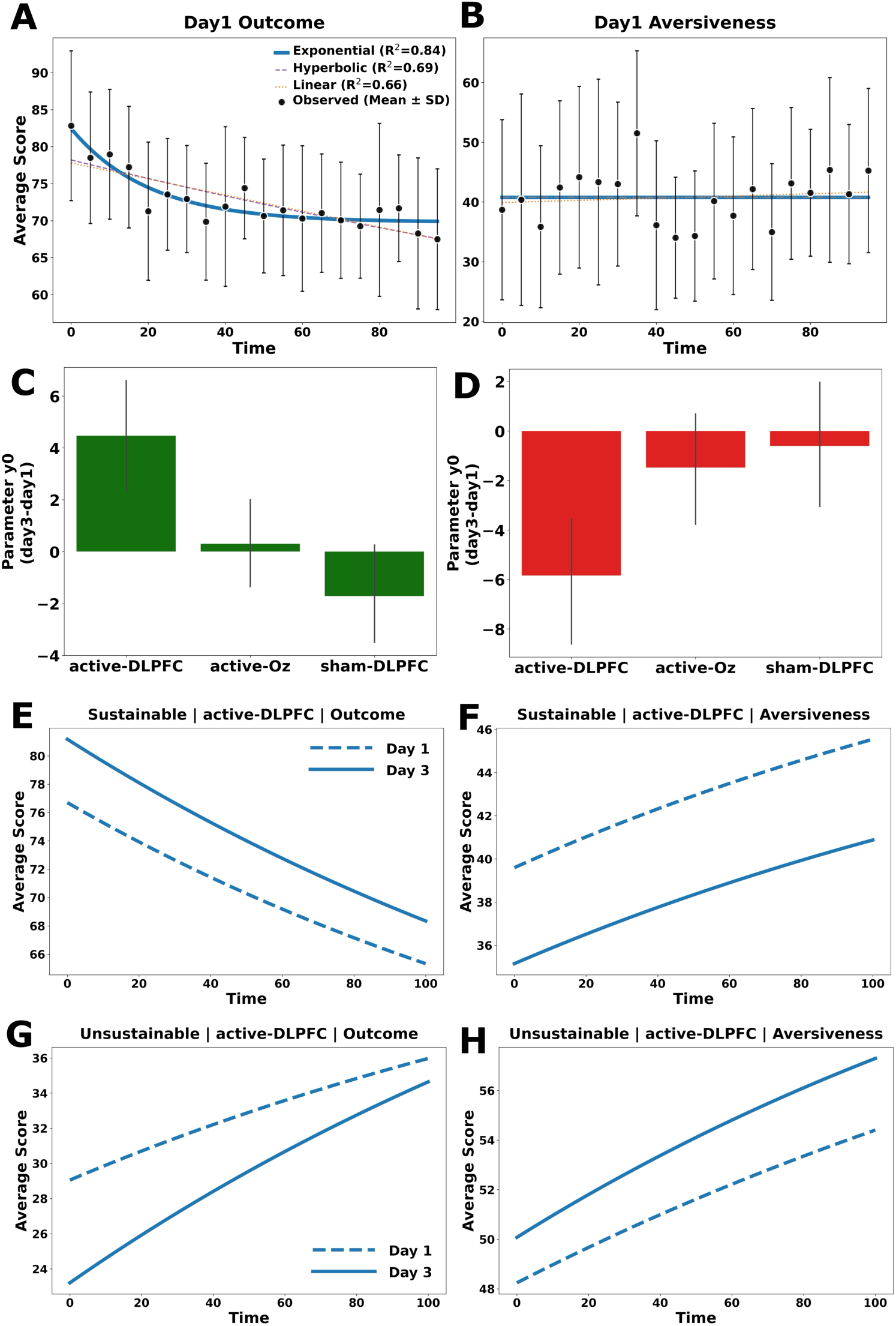
Brain stimulation modulates temporal-value decision model (TVDM) computations underlying sustainable behaviors. **(A)** For sustainable choices, outcome ratings decreased with time and were best captured by an exponential function, outperforming alternative models (hyperbolic, linear, constant).**(B)** By contrast, aversiveness was largely time-invariant and was best captured by a constant model, indicating minimal time-dependent change in aversiveness evaluation. **(C–D)** Change in the outcome-curve intercept (Δy0) by stimulation group. Active DLPFC stimulation increased Δy0 for sustainable choices but decreased Δy0 for unsustainable choices relative to control groups. **(E–H)** Model-implied discounting curves for the active-DLPFC group comparing Day 1 (dashed) versus Day 3 (solid) for sustainable outcome and aversiveness, and unsustainable outcome and aversiveness.

By contrast, for aversiveness, stimulation did not affect any fitted parameters (all omnibus tests ns; *p*’s≥.455 for sustainable, and *p*’s≥.295 for the other condition). Visualizations of model-implied Day 1 vs Day 3 curves in the active-DLPFC group indicate that the most prominent stimulation-related changes were expressed in baseline outcome valuation (**Figure 6E** for sustainable behaviors, **Figure 6G** for unsustainable behaviors), whereas aversiveness curves showed no stimulation-specific parameter shifts (**Figure 6F** and **Figure 6H**). Together, these findings suggest that increasing DLPFC excitability selectively and bidirectionally modulates the baseline (y0) of outcome valuation—enhancing valuation during sustainable decisions while reducing it during unsustainable decisions—without reliable changes in discounting rate or the overall temporal form of aversiveness.

## 3 Discussion

Despite advances in energy and resource efficiency, the large-scale adoption of sustainable technologies and practices ultimately depends on human behaviors and decisions. Here, we integrated computational modelling, behavioral experiments, and non-invasive brain stimulation to identify the neurocomputational mechanisms that constrain sustainable decision-making and to test whether these mechanisms are modifiable. Across two studies, the temporal-value decision model (TVDM) indicated that sustainable decisions are constrained by a temporally structured conflict: sustainable behaviors are experienced as immediately aversive, whereas their outcomes accrue later and are therefore discounted. Critically, anodal HD-tDCS over the lDLPFC increased willingness to choose sustainably, accompanied by higher overall outcome valuation, reduced perceived aversiveness, and a selective upward shift in the outcome-delay function, without reliable changes in the discounting rate or in the temporal parameters governing aversiveness.

### 3.1 Temporal value–aversiveness conflict as a core constraint on sustainable behaviors

The proposed TVDM extends established accounts of intertemporal choice^6,7,53^ by describing a sustainability-relevant asymmetry: outcome values are often delayed and therefore discounted, whereas many sources of aversiveness are experienced immediately. Here, “aversiveness” is operationalized as participants’ subjective cost of acting the depicted behavior at the specified delay, which primarily reflects perceived effort, inconvenience, and momentary unpleasantness of implementation^54,55^. This measure is not intended to capture broader externally imposed costs (e.g., financial expenses, reputational concerns, or norm-based sanctions), which were not manipulated in our paradigm. We focus on this proximal cost signal because models of effortful choice treat subjective effort/inconvenience as a decision-relevant cost that directly competes with anticipated benefits and can discount subjective value at choice^56^.

Model-comparison results further indicated that outcome value alone is neither necessary nor sufficient to explain willingness to act sustainably. Instead, the best-performing account (i.e., Model 3) integrated outcome value and perceived aversiveness and captured individual differences in sustainability through separable sensitivities to each component. This pattern aligns with broader decision frameworks in which choices reflect joint integration of benefits and costs^8,9^ as well as work showing that effort and inconvenience function as decision-relevant costs that can be traded off against anticipated value^57^. Within sustainability research, this offers a computational account of the “knowledge–action gap”: why strong sustainable intentions often fail in the face of minor inconveniences. Specifically, higher perceived environmental values translated into action only when they outweighed immediate aversiveness. However, this condition rarely met because aversiveness is experienced immediately (and thus fully weighted), whereas benefits are delayed and steeply discounted. This interpretation is consistent with neuroeconomic evidence that effort costs can operate as direct discounts on subjective value, distinct from delay discounting^58,59^. By explicitly modelling aversiveness alongside delay, the proposed TVDM extends these frameworks to the environmental decision-making domain.

The “crossover” effect observed in our data—where the value of sustainable outcomes eventually exceeds aversiveness at longer delays or lower effort levels—suggests that a primary barrier to sustainability may not be a lack of concern for the future, but rather the disproportionate weight assigned to immediate physical or psychological effort costs^60^. This parallels the “effort paradox” in cognitive control, whereby individuals value goals but systematically avoid the mental labor required to attain them^60^. The same logic also resembles present-biased procrastination^61^: when costs are immediate, even modest present bias can generate systematic procrastination^22,23,31,49^, whereas the same agent may plan to act when costs are postponed. Together, these parallels suggest that sustainability decisions may share a computational vulnerability with procrastination: future-oriented endorsement coexisting with present-oriented avoidance when the aversive component is immediate.

### 3.2 Integrating TVDM within environmental psychology frameworks

Environmental psychology has developed influential accounts of sustainable behaviors, emphasizing attitudes and norms (for example, TPB)^10,12^, values and moral obligations (VBN)^14^, motivational quality (SDT)^15^, and threat appraisal and coping efficacy (PMT)^17^. Meta-analytic work indicates that these psychosocial determinants reliably relate to pro-environmental behavior, but they do not eliminate substantial unexplained variance and do not fully resolve intention–behavior gaps in contexts where barriers are high^13,62,63^. Integrative reviews similarly highlight that behavior change often requires addressing context, opportunity structures, and the feasibility of action—not merely beliefs^64,65^. By focusing on computational components of decision-making, the TVDM complements intention-based accounts by specifying why sustainable intentions can fail when immediate costs dominate delayed benefits. Concretely, existing frameworks can be mapped onto TVDM components: attitudes, values, and perceived consequences plausibly shape outcome valuation; perceived behavioral control and external constraints plausibly shape aversiveness; and motivational or threat-based processes may influence how these attributes are weighted or retrieved at decision time. This mapping does not imply that the TVDM is a complete theory of sustainable behavior; rather, it offers a computational framework through which diverse determinants may translate into proximal choice computations.

### 3.3 Causal modulation of sustainable decisions by lDLPFC stimulation

A central contribution of the presented brain stimulation study is causal evidence that lDLPFC excitability shapes sustainability-related decisions. Relative to both sham stimulation and an active control montage over occipital cortex, anodal lDLPFC stimulation increased willingness to engage in sustainable behaviors and reduced willingness to engage in unsustainable behaviors. The absence of group-by-delay interactions indicates that this shift was expressed as a broad change across immediate and delayed decisions. These results complement correlational associations linking prefrontal structure and sustainable behaviors^25,34,36,42,66^ and build on mixed prior causal evidence from non-invasive stimulation studies of sustainability^41–43^. Such heterogeneity underscores that stimulation effects may depend on montage, polarity, hemisphere, and task demands^46^, and cautions against simple one-to-one mappings between a stimulation protocol and “more sustainability.” Importantly, our inclusion of both sham and an active-site control increases confidence that the observed effects are not attributable to generic task repetition, somatic stimulation sensations, or non-specific expectancy effects. Nevertheless, the broader tDCS literature also advises caution: quantitative reviews suggest that single-session cognitive effects in healthy populations can be variable and, in some domains, small or inconsistent^67^. The presented results therefore strengthen the case for domain- and task-specific stimulation effects, while also highlighting the importance of independent replication and preregistered confirmatory testing.

### 3.4 What changed under tDCS: valuation level shifts rather than discount-rate shifts

A key contribution of this study is specifying how tDCS altered sustainable decisions. Contrary to the intuitive hypothesis that tDCS might make sustainable behaviors feel “easier” (reduced aversiveness) or attenuate delay sensitivity (lower discounting), our model-based analysis pointed to a different mechanism. tDCS stimulation selectively shifted the baseline outcome valuation parameter, increasing the initial value of sustainable outcomes and decreasing it for unsustainable ones. In contrast, the delay-dependent parameters governing aversiveness showed no stimulation-specific shifts. Together, these patterns indicate that stimulation primarily enhanced baseline outcome valuation rather than altering temporal sensitivity to delay. One possibility is that the lDLPFC supports the resolution of competing decision signals by modulating value representations within valuation circuits. This interpretation aligns with accounts in which the DLPFC integrates attribute information (e.g., environmental impact) into a coherent value signal that influences the ventromedial prefrontal cortex (vmPFC) valuation system^68–70^. By increasing baseline outcome value, tDCS may have enabled delayed environmental benefits to surpass the decision threshold despite the continued presence of immediate effort costs and temporal delay. The stimulation-induced shift in outcome-valuation level—without a corresponding change in discounting rate—therefore fits with a mechanism in which the lDLPFC modulates the weighting or baseline salience of outcomes relevant to sustainability goals, rather than altering the temporal discounting computation itself. Within the TVDM, this pattern can be formalized as an increase in the perceived benefits of sustainable outcomes, thereby biasing choices toward sustainability.

### 3.5 Translational implications for policy and intervention

Beyond mechanistic understanding, the proposed TVDM and presented brain stimulation results have practical implications for intervention design to accelerate behavior change toward global sustainability^71,72^. Interventions should be most effective when they jointly target delayed value and immediate aversiveness. For example, by reducing the effort required for sustainable actions while increasing the salience and credibility of future outcomes. Aversiveness can be reduced through convenient infrastructure, defaults, and choice environments that lower effort and hassle, which may be particularly important because barriers are unevenly distributed across socioeconomic contexts^3,65^. Outcome valuation may be increased by enriching or personalizing future representations (e.g., episodic future simulation)^73–75^ or by emphasizing co-benefits that are less temporally remote than climate outcomes alone.

At the same time, the neuromodulation findings should be interpreted cautiously. We do not view tDCS as an intervention roadmap for population-level climate action. Behavioral interventions targeting household climate action show small average effects and variable persistence across randomized trials^76^, and the broader evidence base indicates that tDCS effects in healthy samples can be inconsistent across tasks and protocols^67^. Moreover, non-invasive brain stimulation raises neuroethical concerns, particularly if framed as a tool for social or moral improvement^77^. Accordingly, the primary value of the presented brain stimulation results lies in using tDCS as a causal probe of the computations that constrain sustainable decisions, which can inform ethically grounded behavioral, educational, and structural interventions rather than replace them. For example, insofar as lDLPFC stimulation facilitated sustainable decisions, the results are consistent with a role for cognitive control processes in overriding immediate (unsustainable) convenience; this motivates testing whether cognitive training approaches that target inhibitory control can indirectly support sustainable behaviors in real-world settings^78^.

### 3.6 Limitations and future studies

Several limitations shape the scope of inference. First, samples comprised healthy young university students, which may limit generalizability across age, cultural, and socioeconomic contexts, particularly given evidence that university populations often report stronger pro-environmental attitudes than the public^79^. To minimize demand characteristics, we have masked the study’s focus on sustainability by presenting it as an investigation of everyday decision-making. Nevertheless, participants selected unsustainable options on a substantial proportion of trials, arguing against a dominant social-desirability account of the observed effects. Second, the primary outcomes were self-report willingness and subjective ratings of aversiveness and outcome value. Although task-based sustainability indices related to a trait questionnaire, self-report can diverge from real-world behavior^80^ and can be influenced by demand characteristics. Future work should incorporate objective behavioral endpoints, ecological momentary assessment, and field-based paradigms to test whether TVDM parameters predict (and can be shifted to change) real-world sustainable actions^81^. Third, the time delay was restricted to days rather than the years-to-decades timescales that dominate many environmental outcomes. Extending the TVDM to longer time delay and incorporating uncertainty, credibility, and collective-action dynamics may be essential for intergenerational contexts^82^. Finally, combining the TVDM with neuroimaging and multivariate predictive modelling could identify neural signatures that track the competing influences of immediate aversiveness and delayed valuation, and test whether these signatures predict interindividual differences in sustainable choice^83^.

### 3.7 Conclusion

Together, our findings suggest that sustainable behavior is shaped not only by knowledge, attitudes, or norms, but also by a computable temporal value–aversiveness trade-off in which immediate task aversiveness competes with discounted future values. By introducing the TVDM and showing that its core computations can be causally perturbed via non-invasive stimulation of the lDLPFC, this work helps explain why sustainable intentions often fail to translate into action, clarifies the computations that govern sustainability-relevant decision, and provides a neuroscience-informed framework for designing ethically grounded interventions to promote sustainable lifestyles across diverse populations.

## Methods

### 1. Participants

In Study 1, we recruited a total of 78 healthy college students. Five participants were excluded for not completing the formal experiment, and three were excluded due to misunderstanding the instructions, resulting in 70 valid participants (mean age 20.39 ± 1.78 years), consisting of 6 males (8.57%) and 64 females (91.43%). In Study 2, the sample size was determined prior to data collection. We conducted a power analysis using GPower 3.1 software^84^ to estimate the required number of participants for detecting reliable effects. (1) Based on the effect size estimated from Study 1 (Cohen’s f = 0.22), a total of 57 participants were deemed necessary to detect significant effects (α= 0.05, β= 0.80, two-way mixed ANOVA). (2) Following conventions from prior mixed within-between-subjects neurostimulation research^48^, which commonly includes 35-40 participants per group, we projected a total recruitment of 120 participants, accounting for potential dropout and discomfort with the neurostimulation. These participants were to be randomly assigned into three groups: active LDLPFC stimulation, sham LDLPFC stimulation, and active Oz stimulation, with approximately 40 participants in each group. Randomization was achieved using block randomization with a block size of 4, which was generated by the online program (randomizer.org). Ultimately, Study2 recruited 133 participants, excluding 5 who did not complete the formal experiment, yielding a final sample of 128 valid participants (mean age 20.50 ± 1.99 years), consisting of 55 males (42.97%) and 73 females (57.03%). Participants were randomly distributed among the three stimulation groups: active LDLPFC stimulation (n = 43; 16 males, 27 females; mean age 21.04 ± 2.20 years), sham LDLPFC stimulation (n = 41; 18 males, 23 females; mean age 21.00 ± 2.17 years), and active Oz stimulation (n = 44; 21 males, 23 females; mean age 19.68 ± 1.44 years). To control for placebo effects, the study used a single-blind design wherein the participants were strictly blinded to their assigned stimulation condition. The efficacy of the blinding procedure was confirmed via participant self-reports (see Supplemental Methods and Results, Table S2, for further details).

Participants in Study 1 and Study2 were screened using identical criteria: all participants were right-handed, had normal or corrected vision, possessed normal intellectual capabilities, and maintained adequate sleep prior to the experiment. They also exhibited a good mental state with sustained attention and had no documented history of psychiatric, physiological, or neurological disorders, no history of pregnancy, no brain injuries, and no contraindications for high-definition transcranial direct current stimulation (HD-tDCS). Informed consent was obtained from each participant prior to the commencement of the experiment. At the conclusion of the study, participants received appropriate compensation. The experimental protocol adhered to the standards of the Declaration of Helsinki and was approved by the Research Ethics Committee of the School of Psychology at Central China Normal University (Wuhan, China) (Ethics Approval No: CCNU-IRB-202312041b)

### 2. Experimental Procedure and Task Paradigm

#### 2.1 Experimental Procedure

In Study 1, participants were asked to perform the sustainable decision-making task (see 2.3 Sustainable decision-making Paradigm below for details). Study 2 followed a two-session design separated by ∼48 hours (Day1-Session 1: baseline; Day3-Session 2: stimulation). On the first session, all three stimulation groups additionally completed the General Ecological Behavior (GEB) questionnaire to assess their sustainable behaviors in daily life. Subsequently, participants completed the same sustainable behavior rating task as in Study 1, without brain stimulation; however, trial timing differed across studies: in Study 1, trials were unlimited, whereas in Study 2, each trial proceeded as follows: a 2-s fixation, a 2-s stimulus description, an aversiveness rating (12 s), an outcome rating (12 s), and a willingness rating (12 s). The order of the aversiveness and outcome questions was counterbalanced across trials. To avoid practice effects, a follow-up experiment was conducted 48 hours later with HD-tDCS. During the follow-up session2, participants were exposed to approximately 30 minutes of either active or sham stimulation. The first 10 minutes required participants to remain at rest, while the final 20 minutes involved doing the formal decision-making task. Finally, participants filled out a questionnaire assessing potential side effects of HD-tDCS. Both the session1 and session2 were conducted between 8:00 AM and 10:00 PM, ensuring that each participant underwent the pre-experiment and follow-up experiment at consistent times.

#### 2.2 Experimental Materials

##### Pictures of Sustainable/non-sustainable behaviors

Stimuli were drawn from Brevers (2021) ^40^ and adapted for a Chinese context by translating and localizing the accompanying descriptions. The repository contains 72 images, consisting of 36 related to sustainable behaviors and 36 related to unsustainable behaviors. However, the depicted behaviors originate from European living contexts, and the accompanying descriptions are in Dutch, making some behaviors challenging for Chinese participants to comprehend. We conducted an independent online validation study (N = 480) in which participants rated each image on clarity and perceived environmental friendliness (See Supplemental Methods and Results; Table S1 for details). Image selection was based on image-level mean and standard deviation ratings (i.e., each image served as one observation in statistical comparisons). The final task used 30 images (15 sustainable, 15 unsustainable) plus 2 practice images available at OSF link listed in Open Science statement below.

##### Questionnaire measure of sustainable behaviors

The General Ecological Behavior (GEB) questionnaire, developed from the studies of Kaiser (1998)^52^ and Kaiser & Wilson (2004)^51^, is recommended in Florian’s (2024)^50^ paper as a standard measure for assessing human pro-environmental behavior. To facilitate responses from Chinese participants while preserving the original meaning, the GEB questionnaire was translated into Chinese by professionals with expertise in English, ensuring complete consistency with the version developed by Kaiser and Wilson (2004)^51^. Detailed information can be found in Supplementary Materials. The questionnaire comprises 50 items, scored using a 5-point Likert scale, with 19 items scored in reverse (item numbers: 4, 5, 8, 9, 12, 14, 15, 16, 21, 25, 27, 29, 30, 31, 32, 33, 35, 40, 41). A higher total score indicates a greater level of pro-environmental behavior among participants.

#### 2.3 Sustainable Decision-Making Paradigm

In the sustainable decision-making paradigm, participants made trial-based decision on certain sustainable/non-sustainable behavior. On each trial, participants viewed an image and brief text describing a particular behavior; critically, no explicit “sustainable/unsustainable” labels were provided. Participants then provided aversiveness/outcome ratings and a willingness rating in sequence. We manipulated temporal delay by varying the time horizon referenced in the questions from the present to 100 days in the future. Each delay constituted a trial lasting 40 s in Study2 (in Study 1, trials were unlimited): 2-s fixation, 2-s description of the sustainable/ unsustainable behavior, 12-s aversiveness or value rating (balanced across trials), and 12-s willingness rating. Each participant completed 30 trials (6 per delay) composed of all materials, with sustainable and unsustainable items (3 each) fully counterbalanced.

##### Task Aversiveness

Task aversiveness denotes the anticipated unpleasantness of performing a given behavior. Participants rated the item “How will you feel using [the above item] or performing [the above behavior] in xx days (selected from the five delays)?” on a 1–100 scale (1 = extremely pleasant, 50 = neutral, 100 = extremely unpleasant). Higher scores indicate greater aversiveness, which has been shown to impede timely action.

##### Outcome Value

Outcome value reflects the perceived future benefit (e.g., environmental improvement) accruing from the behavior. Participants rated the item “Expectation for environmental improvement with [the above item] use or [the above behavior] perform in xx days (selected from the five delays)?” on a 1–100 scale (1 = not at all expected, 50 = uncertain, 100 = extremely expected). Ratings defined the subjective outcome value for each task, with higher scores indicating higher value.

##### Task-Execution Willingness

Task-execution willingness represents the participant’s current intention to perform the behavior. Participants rated the item “Are you willing to use [the above item] or perform [the above behavior] in xx days (selected from the five delays)?” on a 1–100 scale (1 = not willing at all, 50 = uncertain, 100 = completely willing). Higher scores correspond to stronger willingness.

##### Time delay

The time delays started from 0, 5, 10, and 15 days, with subsequent points increasing by 20 days. Delays were drawn from four fixed distributions—(i) 0, 20, 40, 60, 80; (ii) 5, 25, 45, 65, 85; (iii) 10, 30, 50, 70, 90; (iv) 15, 35, 55, 75, 95 days. Each participant was randomly assigned to one of these time distributions and completed the sustainable behavior rating task at the five delays. Within every delay block, the order of aversiveness and value ratings was randomized and counterbalanced across blocks.

### 3. High-Definition Transcranial Direct-Current Stimulation

Our study2 employed the Soterix LTE-tDCS stimulator and the Soterix 4×1 multi-channel stimulation adapter for high-definition transcranial direct current stimulation (HD-tDCS) (Soterix Medical Inc., New York, NY). HD-tDCS used a 4×1 ring electrode configuration, with the central electrode positioned over the target area and surrounded by four return electrodes. Given the nonlinear relationship between electrode resistance and the electrochemical processes at the electrode-skin interface^85^, we measured impedance in “quality units” for the HD-tDCS. Based on prior research, each electrode’s quality unit was maintained at or below 1.5 to 2.0^86,87^. For anodal stimulation of the left dorsolateral prefrontal cortex (LDLPFC), the central electrode was positioned at F3 according to the International 10-20 electrode placement system. The four return electrodes were respectively placed at F5, AF3, FC3, and F1^88^. For anodal stimulation of the occipital cortex, the anode was placed at Oz^89^, with corresponding return electrodes at POz, O1, O2, and Iz.

Prior to stimulation, we separated the hair beneath the electrodes to expose the scalp and injected approximately 2 mL of conductive gel into the electrode housing. Numerous studies have indicated that a current intensity of 1.5 mA is safe for healthy participants and enhances brain function^90–93^. Specifically, during active stimulation, a constant 1.5 mA current was maintained for 30 minutes, which included an initial 30 seconds of gradual current increase and a final 30 seconds of gradual decrease. In the sham stimulation condition, the current decreased from 1.5 mA to 0 over 30 seconds, with the electrodes remaining in position for 29 minutes and 30 seconds. This setup ensured that participants were blinded to the stimulation condition through physical sensations on the scalp without influencing neural function^94,95^. The choice of these sham stimulation parameters was based on previous studies indicating that skin sensations caused by tDCS (e.g., tingling) typically diminish within the first 30 seconds of stimulation^96,97^. Moreover, studies have demonstrated that participants receiving active stimulation for only 30 seconds reported discomfort levels, sensation duration, attention, and fatigue comparable to those in the sham group. Multiple studies have also shown that this sham stimulation has no effect on various cognitive processes, including working memory, motor performance, impulse control, and emotional regulation^90–93^.

In Study 2, participants were instructed to avoid checking their phones and to relax or remain in a resting state during the initial 10 minutes of stimulation, followed by a 20-minute continuous stimulation while completing the rating task, to mitigate potential confounding effects. All participants completed the “tDCS Adverse Effect Assessment Questionnaire” before and after stimulation, which consisted of three sections: (a) changes in emotional state, (b) perceived stimulation intensity, and (c) ten common potential adverse reactions following tDCS (e.g., headaches, neck pain, scalp pain, numbness, itching)^98^. Participants did not report any adverse reactions, experiencing only mild skin tingling during the current variations (See Supplemental Methods and Results; Table S2 for details).

### 4. Data and Statistical Analyses

#### 4.1 Overview of datasets and variables

Analyses were conducted on trial-level behavioral data from the sustainable decision-making task. Each trial included (A) a stimulus identifier (a participant sustainable or unsustainable behavior), (B) participant identifier (subject), (C) a temporal delay (time; 0–100 scale), and three ratings collected on 1–100 scales: perceived aversiveness (aversiveness), perceived outcome value (outcome), and willingness to perform the behavior (willingness). All analyses were performed separately for sustainable and unsustainable trials when appropriate. Data from Study1 and Study2 were analyzed identically, except for the comparisons between stimulations

#### 4.2 Temporal trend analyses (delay effects)

To quantify whether ratings changed systematically with temporal delay, we estimated within-participant linear trends across different time delay, separately for sustainable and unsustainable trials. For each condition and each metric (aversiveness, outcome, willingness), we firstly averaged ratings within each participant and delay level (subject × time) to avoid overweighting repeated trials at the same delay. For each participant, fit a linear model using least-squares slope estimation:

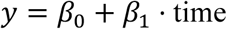

where 𝛽_1_ is the participant-specific temporal slope.

We tested whether slopes differed from zero at the group level using a one-sample t-test on participant slopes.

#### 4.3 Computational modelling of willingness using outcome and aversiveness

##### Candidate models

To test whether willingness is best explained by perceived outcome value, perceived aversiveness, or their combination, we evaluated four candidate models predicting trial-wise willingness. All models were fit separately for sustainable and unsustainable trials.

- **Model 1 (Outcome-only):**

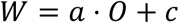

- **Model 2 (Aversiveness-only):**

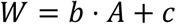

- **Model 3 (Additive outcome + aversiveness):**

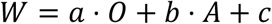

- **Model 4 (Additive + interaction):**

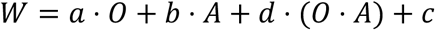

Here, 𝑊 denotes willingness, 𝑂 outcome value, 𝐴 aversiveness, 𝑐 an intercept term, and 𝑑 the interaction coefficient.

##### Estimation procedure

Models were fit using ordinary least squares (OLS) with an assumed Gaussian observation model. For each fit, we computed: (1) fitted values and residuals, (2) Gaussian log-likelihood at the maximum-likelihood estimate of 𝜎^2^(using SSE/n), and (3) in-sample 𝑅^2^.

##### Model comparison

We compared models using three complementary approaches: (1) Information criteria (AIC/BIC). AIC and BIC were computed from the Gaussian log-likelihood. Parameter counting included regression coefficients plus one variance parameter 𝜎^2^. (2) Leave-one-out cross-validation (LOO-CV) using exact linear-regression LOO. Pointwise LOO predictive log densities were computed from analytic leave-one-out predictions using hat-matrix diagonals. Summed pointwise values yielded ELPD_LOO_ for each model. We additionally computed pairwise ΔELPD values relative to a reference model (Model 3) and estimated uncertainty from pointwise differences:

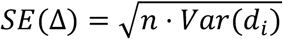

where 𝑑_$_is the pointwise difference in LOO log density between the reference model and the comparison model.

##### Grouped k-fold cross-validation by participant

To ensure evaluation generalized across individuals, we used GroupKFold cross-validation with grouping by participant (subject). For each fold, models were refit on the training data and evaluated on held-out trials. We summarized predictive performance by: held-out predictive log-likelihood (sum of Gaussian log densities under the training-fit variance), and mean squared error (MSE) on held-out data.

##### Inference for Model 3 coefficients

For interpretability and hypothesis testing, we estimated regression coefficients for Model 3 using OLS with cluster-robust standard errors clustered by participant to account for within-subject dependence. We then assessed the unique contribution of each predictor using robust Wald tests in the full Model 3 fit (testing 𝑎 = 0and 𝑏 = 0 separately). When fitting Model 4 for completeness, the interaction term was tested using the same cluster-robust procedure.

##### Individual-level parameter estimation and condition comparisons

To obtain participant-specific parameter estimates for Model 3, we fit Model 3 separately to each participant’s trials within each condition using least-squares curve fitting:

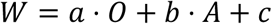

This produced per-participant estimates of outcome sensitivity (𝑎), aversiveness sensitivity (𝑏), and intercept (𝑐) for sustainable and unsustainable trials. To compare parameters between sustainable and unsustainable conditions, we performed **paired-sample t-tests** across participants for each parameter (𝑎 and 𝑏)

#### 4.3 Outcome–aversiveness dynamics by sustainable decisions

To test whether sustainable decisions occur preferentially when perceived outcomes outweigh aversiveness, we discretized willingness into four ordinal decision categories using fixed bins on the 1–100 scale: not to do (1–25); reluctant (26–50); consider (51–75); to do (76–100). Within each condition (sustainable / unsustainable), we computed subject-level mean outcome and aversiveness within each decision category. We tested whether the relationship between metric (outcome vs weighted aversiveness) and decision category depended on their interaction using a 2 (metric) × 4 (decision category) repeated-measures ANOVA implemented with repeated ANOVA.

##### Net value analysis (Outcome − Aversiveness)

We defined a net value signal within each category:

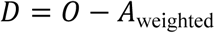

and tested whether 𝐷 varied across decision categories using a one-way repeated-measures ANOVA on 𝐷. We additionally tested a linear trend across the ordered categories.

#### 4.4 Hyperbolic discounting analyses and crossover dynamics

##### Hyperbolic model

To characterize temporal dynamics of outcome and aversiveness ratings, we fit a hyperbolic function:

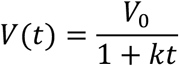

where 𝑉_!_captures the estimated value at 𝑡 = 0 and 𝑘 captures the discounting rate.

Fits were performed using nonlinear least-squares (curve_fit) with parameter bounds constrained to nonnegative values (𝑉_!_ ≥ 0, 𝑘 ≥ 0).

##### Participant-level discounting fits

For each participant and condition, we fit separate hyperbolic functions to: outcome values across time; aversiveness values across time. These participant-level estimates were used to test within-condition asymmetries in discounting rates (e.g., whether aversiveness declines faster than outcome) using paired t-tests and (as a robustness check) Wilcoxon signed-rank tests. Between-condition comparisons of discounting parameters were evaluated using Welch’s independent-samples t-tests.

##### Crossover time between outcome and aversiveness

To quantify when outcome overtakes aversiveness across time, we computed the earliest time point 𝑡 satisfying:

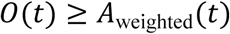

using a dense grid search over the observed horizon (0–100 scale; step size 0.05). Crossover was computed both for: group-level curves (using group-mean fitted parameters), and participant-level fits, summarized as the proportion of participants exhibiting a crossover within the horizon and the median crossover time among those who crossed.

##### Decision distribution dynamics across time

To link temporal dynamics to decision outcomes, we examined how the proportions of categorical responses changed across delay. We focused on the extreme categories “to do” **and** “not to do”. For each delay level and condition, we computed the percentage of trials labeled as “to do” and “not to do,” then fit hyperbolic functions of delay to these percentage trajectories using the same form as above. Because these fits were performed at the condition level (single curve per condition), parameters were reported descriptively rather than subjected to inferential testing.

##### Discounting trajectories conditioned on decision outcome

To dissociate valuation dynamics for “to do” vs “not to do” decisions, we refit hyperbolic trajectories separately within each decision outcome subset. Specifically, within each condition we fit hyperbolic functions to outcome and aversiveness ratings as functions of delay for trials ending in “to do” and trials ending in “not to do”.

#### 4.5 tDCS effects on underlying TVDM computations

To identify computational mechanisms through which anodal tDCS over DLPFC altered sustainable decision-making, we quantified (A) changes in evaluative components (Outcome and Aversiveness ratings) and (B) changes in model-derived parameters from Day 1 (baseline) to Day 3 (stimulation session). Analyses were conducted separately for sustainable and unsustainable trials and compared across stimulation groups. All analyses were implemented in Python using standard scientific libraries (NumPy, pandas, SciPy, scikit-learn). Model fitting used nonlinear least squares (SciPy curve_fit and least_squares). Unless otherwise noted, statistical tests were two-tailed.

##### tDCS effects on evaluative components (Outcome and Aversiveness)

To test whether stimulation altered evaluative components supporting behavioral change, we focused on within-participant change from Day 1 to Day 3 for the relevant rating measures (Outcome and Aversiveness), computed as: Δ𝑋=𝑋_Day 3_−𝑋_Day 1_. These change scores were then compared across groups using one-way between-subjects ANOVA

##### tDCS effects on Model 3 parameters

To test whether stimulation altered the mapping from evaluations to willingness, we fit Model 3 (see above for details of Model3 fitting) separately for sustainable and unsustainable trials and separately for Day 1 and Day 3. In model3, where a captures outcome sensitivity, b captures aversiveness sensitivity, and c is an intercept-like constant. Parameters were estimated individually for each participant using nonlinear least squares on trial-level data. For each participant and condition (i.e., sustainable/unsustainable), we computed parameter change scores and were compared across stimulation group.

##### Temporal functional-form comparison

To characterize the temporal form of ratings as a function of delay, we compared candidate models fit to the mean time series across participants. For each metric (Outcome and Aversiveness), we first computed participant-averaged values per timepoint, then computed the grand mean and standard deviation at each timepoint.

We then fit four candidate functions to the mean time series:

1. **Constant (null):** 𝑦(𝑡) = 𝑐
2. **Linear:** 𝑦(𝑡) = 𝑚𝑡 + 𝑏
3. **Generalized hyperbolic:** 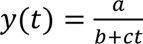
4. **Generalized exponential:** 𝑦(𝑡) = 𝑎exp (𝑏𝑡) + 𝑐

Model fits were compared using AIC, BIC, and 𝑅^2^. Akaike weights were computed from AIC differences:

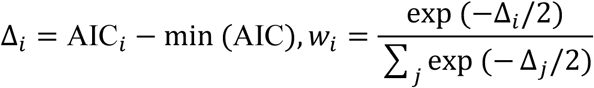

The best-supported functional form for each metric was selected based on AIC (and convergent evidence from BIC/weights), matching the model-selection results reported in the Results section.

##### Individual temporal fits using the winning functional form

Using the winning temporal form, we fit each participant’s delay-dependent ratings separately for sustainable and unsustainable trials, and separately for Day 1 and Day 3. Individual fits used a numerically stable exponential-decay parameterization:

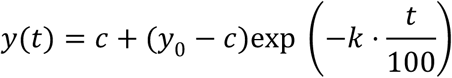

where:

- *y*_0_ is an intercept-like baseline level (value at minimal delay),
- is the asymptote/offset,
- is the decay rate.

##### Fitting procedure

For each participant, trials were first averaged within each unique delay value to reduce trial noise. Parameters were estimated with constrained nonlinear least squares. Rating bounds were constrained to the scale of the dependent variable (0–100). The decay rate was constrained to be non-negative and capped such that the function retained at least half its amplitude across the full delay range (implemented as 𝑘 ≤ ln 2). Robust fitting used a soft L1 loss to reduce the influence of outliers. Model fit quality was summarized using mean squared error (MSE) and 𝑅^2^.

For compatibility with the generalized exponential form reported in some plots/outputs, parameters were also expressed as:

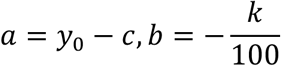

Change scores were computed for the parameters (e.g., Δ𝑦_!_, Δ𝑘, Δ𝑐) as Day 3 minus Day 1 and tested across stimulation groups.

## 4.6 Data availability

The raw data from the experiments as well as the data displayed in each figure are available via https://osf.io/2bxvq/overview?view_only=3f51bd0d4ff24c38901faaecf3553533

## 4.7 Code availability

The code for running all experiments and analyses and for generating the figures is available via https://osf.io/2bxvq/overview?view_only=3f51bd0d4ff24c38901faaecf3553533

## Acknowledgements

This work was supported by the National Natural Science Foundation of China (grant No. 32300879), Humanities and Social Sciences Fund, Ministry of Education (grant No. 22YJCZH109). The funders had no role in the study design, data collection and analysis, decision to publish or preparation of the manuscript.

## Author contributions

W.L. conceived the study. W.L. and X.Y.Z. designed the experiment. X.Y.Z. implemented and conducted the experiments. W.L. and X.Y.Z. analyzed the data. W.L. and X.Y.Z. wrote the manuscript.

