## Supplemental files for "Non-invasive brain stimulation biases temporal value–aversiveness computations and promotes sustainable decision-making"

**for**

**This file contains**

Supplemental Methods and Results

Supplemental Figures S1-11

Supplemental Tables S1-2

Supplemental Questionnaires

*** Correspondence**:

Dr. Wei Liu, School of Psychology, Central China Normal University

Address: the 8th floor, Nanhu Complex Building, No 152 Luoyu Road, Wuhan.

Postal Code: 430079

### Supplemental Methods and Results

#### Overview


1. Validation and selection of sustainability image stimuli
2. Computational model comparison methods and results
3. Weighted averiveness and outcome for comparsion
4. Replication analyses of Study2
5. tDCS safety, tolerability, and blinding checks
6. Delay-Invariance of tDCS Effects on Outcome and Aversiveness
7. Effects of tDCS on Computational Parameters (Model 3)


#### 1. Validation and selection of sustainability image stimuli

To evaluate and select experimental materials (i.e., pictures and explanatory texts), we recruited participants through an online survey platform (https://www.wjx.cn). We obtained 480 valid adult responses (256 men, 224 women; mean age = 21.51 ± 1.98 years) from 29 provincial-level administrative regions in mainland China. Participants were compensated for completing the Sustainable Materials Online Evaluation Questionnaire (mean completion time = 1,056.43 s). Participants rated each of 72 images on five dimensions using a 7-point Likert scale (1–7; higher scores indicate greater agreement): (Q1) whether the depicted item/behavior is commonly used in China; (Q2) whether it is commonly encountered in daily life; (Q3) understanding of its purpose or use; (Q4) ease of classifying it as environmentally friendly versus unfriendly; and (Q5) perceived environmental friendliness.

To quantify overall comprehensibility, we aggregated scores from Q1–Q4 into a single index. We additionally examined Q4 and Q5 as indicators of (i) the ease of determining environmental friendliness and (ii) suitability for environmentally focused research (Table S1).

A two-way between-subjects ANOVA on the questionnaire data showed that the selected materials (32 images; 30 for task; 2 for practice) scored higher on the comprehensibility items (Q1–Q4) than the unselected materials, with lower variability across items (means: $F(1, 68) = 4.26$, $p = 0.04$, $\eta^2 = 0.05$; standard deviations: $F(1, 68) = 4.36$, $p = 0.04$, $\eta^2 = 0.05$). Thus, the selected images were more closely aligned with participants' lived experience and easier to understand. The selected materials also yielded higher Q4 scores than the unselected materials, again with greater consensus across items (means: $F(1, 68) = 26.52$, $p < 0.001$, $\eta^2 = 0.26$; standard deviations: $F(1, 68) = 24.35$, $p < 0.001$, $\eta^2 = 0.25$), indicating that they were easier to classify as environmentally friendly versus unfriendly. Consistent with this categorization, Q5 ratings indicated that sustainable images were judged as more environmentally friendly than unsustainable images both across the full set of 72 images ($t(70) = 11.90$, $p < 0.001$, Cohen's $d = 2.80$) and within the selected set ($t(70) = 13.11$, $p < 0.001$, Cohen's $d = 4.64$). Finally, a 2 (sustainability category: sustainable vs unsustainable) × 2 (selection status: selected vs unselected) ANOVA on Q5 ratings across all materials revealed a significant interaction (means: $F(1, 68) = 26.96$, $p < 0.001$, $\eta^2 = 0.094$; standard deviations: $F(1, 68) = 26.47$, $p < 0.001$, $\eta^2 = 0.103$), indicating that the separation between sustainable and unsustainable categories was larger for the selected materials but attenuated for the unselected materials. Together, these results support the use of the selected image set for eliciting reliable judgements of environmental friendliness in sustainability-focused research.

#### 2. Computational model comparison methods and results

##### 2.1 Overview and goal

Using trial-by-trial data from Study 1, we evaluated candidate computational models of participants' willingness to perform each type of behavior (**Figure S2**). Our primary aim was to test whether perceived outcome value and task aversiveness jointly predict willingness, and whether adding an outcome-by-aversiveness interaction improves predictive performance. Model comparison was conducted separately for sustainable and unsustainable trials, consistent with the main analyses.

##### 2.2 Data and predictors

Each trial included:

- **Willingness** rating (dependent variable; continuous)
- **Outcome** rating (predictor; perceived benefit/value)
- **Aversiveness** rating (predictor; perceived cost/effort)

To avoid dependence artifacts from repeated trials within individuals, all regression-based inferential tests used cluster-robust (subject-clustered) standard errors. Predictive evaluation used subject-wise cross-validation where specified.

##### 2.3 Candidate models

Let $y$ denote willingness, $O$ denote outcome, and $A$ denote aversiveness. We compared four models:

- **Model 1 (Outcome-only):**

$$y = \beta_O O + \beta_0$$

- **Model 2 (Aversiveness-only):**

$$y = \beta_A A + \beta_0$$

- **Model 3 (Additive integration):**

$$y = \beta_O O + \beta_A A + \beta_0$$

- **Model 4 (Interaction):**

$$y = \beta_O O + \beta_A A + \beta_{OA}(O \times A) + \beta_0$$

All models were fit as Gaussian linear models using ordinary least squares (OLS). Because Models 1–4 are linear in parameters, OLS provides the exact maximum-likelihood estimates under Gaussian residual assumptions.

Model comparison used LOO-CV ELPD and information criteria; ΔELPD was defined as ($\Delta ELPD = ELPD_{Model3} - ELPD_{comparison}$), so positive values favor Model 3. Subject-wise generalization was assessed using 5-fold GroupKFold by participant.

##### 2.4 Model evaluation metrics

We assessed model performance using complementary indices that balance fit and complexity:

1. **Leave-one-out cross-validation (LOO-CV):**

   We computed the expected log predictive density for each model ($ELPD_{LOO}$), and report model differences as:

$$\Delta \text{ELPD} = \text{ELPD}_{\text{Model 3}} - \text{ELPD}_{\text{comparison}}$$

Standard errors for ΔELPD were computed from pointwise LOO log predictive densities, following standard LOO practice.

2. **Information criteria (AIC/BIC):**

   AIC and BIC were computed from the Gaussian log-likelihood with maximum-likelihood residual variance. Lower values indicate better tradeoff between fit and complexity.

3. **Subject-wise cross-validated predictive performance (GroupKFold):**

   To evaluate generalization to new participants, we performed 5-fold GroupKFold cross-validation by subject, ensuring all trials from a participant appeared in either training or test but not both. For each fold, we computed:

- Cross-validated predictive log-likelihood (sum of log densities on held-out trials)
- Cross-validated mean squared error (CV-MSE; avg_mse)

This combination allowed us to assess both likelihood-based predictive accuracy and error-based performance.

#### 2.5 Inferential evidence for "joint prediction"

In addition to model selection, we tested whether outcome and aversiveness each explained unique variance in willingness using Model 3 with subject-clustered standard errors. We report coefficient estimates with 95% confidence intervals and p-values, and we conducted robust Wald tests for nested contributions. We report:

- coefficient estimates with 95% CIs and p-values for Model 3
- robust Wald tests evaluating whether each predictor contributes beyond the other:
    - $H_0: \beta_A = 0$ (aversiveness adds beyond outcome)
        - $H_0: \beta_O = 0$(outcome adds beyond aversiveness)

#### 2.6 Supplementary Model Comparison Results

##### 2.6.1 Sustainable Context

**Model comparison.** Across metrics, Model 3 (outcome + aversiveness) provided the best balance of predictive accuracy and parsimony. Model 3 yielded the lowest AIC and the best LOO-CV performance among candidate models (ELPD_LOO = −4579.83; AIC = 9152.78; BIC = 9172.01; CV log-likelihood = −4589.67; CV-MSE = 363.60; $R^2$ = 0.544). Relative to Model 3, both single-predictor models performed substantially worse (Model 1: ΔELPD = 323.06, SE = 35.39; Model 2: ΔELPD = 28.77, SE = 8.84). Adding an interaction term did not meaningfully improve out-of-sample prediction (Model 3 vs. Model 4: ΔELPD = 1.61, SE = 0.85) and slightly worsened AIC (Model 4 AIC = 9154.32).

**Coefficient evidence (cluster-robust).** In Model 3, both predictors were significant in the expected directions: outcome positively predicted willingness (β = 0.237, 95% CI [0.141, 0.333], p < .001) and aversiveness negatively predicted willingness (β = −0.699, 95% CI

[−0.779, −0.619], $p < .001$). Robust nested contribution tests indicated that each predictor explained unique variance beyond the other (aversiveness beyond outcome: $\chi^2 = 295.28$, $p < .001$; outcome beyond aversiveness: $\chi^2 = 23.58$, $p = 0.000001$). Consistent with these results, the interaction term in Model 4 was not significant ($\beta\_OA = -0.0007$, $p = .656$).

**Conclusion (sustainable).** Sustainable willingness is best captured by an additive integration of outcome value and aversiveness (Model 3), with no evidence that an interaction is required.

##### 2.6.2 Unsustainable trials

**Model comparison.** Unsustainable trials showed the same qualitative pattern—both predictors contributed—while the interaction model showed limited and uncertain incremental benefit. Model 3 strongly outperformed the outcome-only model (Model 1: ΔELPD = 313.70, SE = 32.00) and modestly outperformed the aversiveness-only model (Model 2: ΔELPD = 5.82, SE = 4.26). In information-criterion terms, Model 4 achieved the lowest AIC/BIC (Model 4 AIC = 9171.37) relative to Model 3 (AIC = 9180.95). However, the LOO-CV difference between Models 3 and 4 was small and imprecise (Model 3 vs. Model 4: ΔELPD = −4.23, SE = 4.19), indicating that the additional flexibility in Model 4 yields, at most, a modest and not clearly reliable gain in out-of-sample predictive accuracy.

**Coefficient evidence (cluster-robust).** In Model 3, both predictors were significant: outcome positively predicted willingness ($\beta = 0.109$, 95% CI [0.016, 0.202], $p = .022$) and aversiveness negatively predicted willingness ($\beta = -0.698$, 95% CI [−0.779, −0.617], $p < .001$). Robust nested contribution tests again supported unique contributions of both predictors (aversiveness beyond outcome: $\chi^2 = 286.83$, $p < .001$; outcome beyond aversiveness: $\chi^2 = 5.28$, $p = .0216$). In Model 4, the interaction term was significant ($\beta\_OA = 0.0034$, $p = .015$), suggesting some coupling between outcome and aversiveness for unsustainable willingness. Notably, the main effects in Model 4 were less stable than in

Model 3 (with outcome becoming non-significant when the interaction was included), consistent with the small and uncertain incremental predictive advantage under LOO-CV.

**Conclusion (unsustainable).** Although Model 4 marginally improved information criteria and produced a significant interaction term, its predictive advantage over Model 3 was small and not clearly reliable under LOO-CV. Across both trial types, the robust and consistent finding is that outcome value and aversiveness jointly predict willingness, which is captured parsimoniously by Model 3.

##### 3. Weighted aversiveness and outcome for comparison

We found that task aversiveness exerted a stronger influence on sustainable decisions than outcome value. To express outcome value and aversiveness on a common scale, we estimated the optimal scaling weight for aversiveness relative to outcome value and found that a weight of 1.44 maximized model performance. Accordingly, we scaled aversiveness ratings by 1.44 in subsequent analyses to directly compare the effects of aversiveness and outcome value on choice.

Specifically, we estimated an empirical scaling weight for aversiveness relative to outcome value: we searched for the weight $w$ that best aligned a single composite predictor with participants' trial-wise willingness ratings. For each candidate weight $w \in [0,4]$ (step = 0.001), we constructed a composite decision variable

$$x(w) = \text{Outcome} - w \cdot \text{Aversiveness},$$

and fit an ordinary least squares model:

$$\text{Willingness} = \beta\, x(w) + \alpha.$$

We selected the value of $w$ that minimized mean squared error (equivalently maximizing $R^2$ given the same outcome variable). This procedure yielded an optimal scaling factor of $w = 1.44$, indicating that a one-unit increase in rated aversiveness corresponded to an $\sim 1.44$-unit decrease in outcome value in terms of its association with willingness (**Figure S4**). At this optimal weight, the fitted model parameters were $\beta = 0.0895$ and $\alpha = 87.1638$, with model performance $\text{MSE} = 367.93$ and $R^2 = 0.5069$.

Accordingly, in subsequent analyses we rescaled aversiveness ratings by this factor ($\text{Aversiveness}_{\text{scaled}} = 1.44 \times \text{Aversiveness}$) to directly compare the effects of aversiveness and outcome value on choice on the same scale.

#### 4. Replication analyses of Study2

##### 4.1 Computational Modeling of Willingness

***Methods:*** To evaluate the predictive utility of perceived outcome value and task aversiveness on participants' willingness to engage in sustainable and unsustainable behaviors, we compared four candidate computational models using trial-by-trial data. Model 1 and Model 2 assumed willingness was driven solely by outcome value or task aversiveness, respectively. Model 3 integrated both factors additively, while Model 4 included their interaction. Model performance was assessed via leave-one-out cross-validation (LOO-CV; evaluating changes in expected log predictive density, $\Delta\text{ELPD}$), Akaike and Bayesian Information Criteria (AIC/BIC), and participant-wise cross-validated predictive log-likelihood (GroupKFold).

***Results:*** For sustainable trials, the additive model (Model 3) provided the best fit. It significantly outperformed both the outcome-only model ($\Delta\text{ELPD} = 649.48,\ \text{SE} = 47.84$) and the aversiveness-only model ($\Delta\text{ELPD} = 19.61,\ \text{SE} = 7.47$), while also yielding the lowest AIC (15,930.63). Cluster-robust regression confirmed that outcome value positively predicted willingness ($\beta = 0.182,\ p < .001$), whereas task aversiveness was a negative predictor ($\beta = -0.720,\ p < .001$). Both parameters accounted for unique variance (robust Wald tests: $\chi^2 = 22.42,\ p < .001$ for outcome; $\chi^2 = 437.46,\ p < .001$ for aversiveness). Incorporating an interaction term (Model 4) did not improve predictive performance ($\beta =$

$-0.0004,\ p = .784$; $\Delta\text{ELPD} = 1.65,\ \text{SE} = 0.53$relative to Model 3).

For unsustainable trials, model comparisons yielded largely convergent results. LOO-CV supported Model 3 over the single-factor models (Model 1: $\Delta\text{ELPD} = 319.20,\ \text{SE} = 230.96$; Model 2: $\Delta\text{ELPD} = 22.16,\ \text{SE} = 8.73$) and the interaction model ($\Delta\text{ELPD} = 63.30,\ \text{SE} = 45.04$). Although AIC marginally favored Model 4 ($\text{AIC} = 16{,}427.74$), the interaction term was not statistically reliable ($\beta = -0.0083,\ p = .113$), indicating negligible incremental benefit. Within Model 3, outcome value ($\beta = 0.187,\ p < .001$) and aversiveness ($\beta = -0.572,\ p < .001$) remained significant independent predictors. Consistent with Study 1, Model 3 was retained as the primary computational account, robustly capturing the joint additive contributions of outcome value and task aversiveness across all trial types.

##### 4.2 Categorical Integration of Outcome and Aversiveness

***Methods:*** To elucidate how participants integrate perceived outcomes and task aversiveness—specifically testing whether sustainable actions are most probable when perceived outcomes outweigh costs—we stratified trials into four decision categories based on willingness ratings: "not to do", "reluctant", "consider", and "to do". We quantified the absolute outcome and aversiveness ratings within each category, alongside their relative balance (net value, $D = \text{Outcome} - \text{Aversiveness}$). Data were analyzed using repeated-measures ANOVAs to assess the Metric (Outcome vs. Aversiveness) $\times$Decision Category interaction.

***Results:*** Data from sustainable trials ($N = 49$ complete cases; 79 excluded due to missing cells) supported the proposed integration account. A 2 $\times$4 repeated-measures ANOVA revealed a significant Metric $\times$Decision Category interaction ($F(3{,}144) = 75.00,\ p < .001$, $\eta_p^2 = .610$; **Figure S9A**). Holm-corrected paired t-tests indicated that for low-willingness categories, perceived aversiveness significantly exceeded outcome value ("not to do":

$M_{\text{diff}} = -27.93,\ t(59) = -7.19,\ p < .001,\ d_z = -0.93$; "reluctant": $M_{\text{diff}} = -15.17$, $t(103) = -7.42,\ p < .001,\ d_z = -0.73$). This balance inverted as willingness increased: outcome value surpassed aversiveness in the "consider" category ($M_{\text{diff}} = 12.84,\ t(125) = 7.89,\ p < .001,\ d_z = 0.70$) and exhibited a substantial advantage in the "to do" category ($M_{\text{diff}} = 36.78,\ t(120) = 22.74,\ p < .001,\ d_z = 2.07$). Consequently, net value $D$ demonstrated a reliable within-subject monotonic increase across categories ($F(3,144) = 75.00,\ p < .001,\ \eta_p^2 = .610$), highlighted by a strong linear trend ($t(48) = 12.90,\ p < .001,\ d_z = 1.84$). This reflects a systematic transition from negative to positive net value as participants shifted from refusal to action.

Unsustainable trials ($N = 71$ complete cases; 57 excluded) exhibited a distinctly different integration profile. The repeated-measures ANOVA showed a robust Metric ×Decision Category interaction ($F(3,210) = 153.74,\ p < .001,\ \eta_p^2 = .687$; **Figure S9B**). Although outcome value increased ($M = 23.16$ to $36.79$) and aversiveness decreased ($M = 109.49$ to $45.95$) with greater willingness, aversiveness consistently exceeded outcome value across all categories. Planned comparisons confirmed negative net values universally: "not to do" ($M_{\text{diff}} = -86.33,\ t(101) = -36.74,\ p < .001,\ d_z = -3.64$), "reluctant" ($M_{\text{diff}} = -58.28$, $t(123) = -34.29,\ p < .001,\ d_z = -3.08$), "consider" ($M_{\text{diff}} = -32.31,\ t(121) = -19.79$, $p < .001,\ d_z = -1.79$), and "to do" ($M_{\text{diff}} = -9.16,\ t(93) = -3.23,\ p = .002,\ d_z = -0.33$). While net value became less negative at higher willingness levels (linear trend: $t(70) = 17.13,\ p < .001,\ d_z = 2.03$), it failed to cross into positive territory. This indicates that increased willingness to execute unsustainable actions is not driven by the subjective dominance of outcomes over costs.

#### 5. tDCS safety, tolerability, and blinding checks (Study 2)

We analyzed each of the three sections of the tDCS Adverse Effect Assessment

Questionnaire separately (**Table S2**). Overall, participants reported no adverse reactions, noting only mild skin tingling during current changes. Self-reported emotional state (pleasantness/activation; 1–9) did not differ across the three groups before versus after tDCS in Study 2 ($F(2, 125) = 1.40$, $p = 0.25$, $\eta^2 = 0.02$). Within-group comparisons likewise showed no significant pre–post changes (active LDLPFC: $t(125) = 1.13$, $p = 1.00$, $d = 0.16$; sham LDLPFC: $t(125) = 1.33$, $p = 1.00$, $d = 0.19$; active Oz: $t(125) = 2.24$, $p = 0.40$, $d = 0.30$). These results indicate that real or sham tDCS did not measurably alter participants' emotional state, suggesting that group differences observed in the Sustainable Behavior Rating Task were unlikely to be driven by emotion changes.

Perceived stimulation intensity (1–9) did not differ between the active and sham LDLPFC groups ($t(120) = 2.50$, $p = 0.05$, $d = 0.56$), between the active LDLPFC and active Oz groups ($t(120) = 0.76$, $p = 0.75$, $d = 0.17$), or between the active Oz and sham LDLPFC groups ($t(120) = 1.77$, $p = 0.21$, $d = 0.39$). Thus, participants were unable to reliably distinguish stimulation conditions, and the task effects were unlikely to reflect expectancy or placebo influences. Finally, ratings of adverse symptoms after stimulation (headache, neck/scalp pain, numbness, itching, burning, skin redness, drowsiness, difficulty concentrating, or acute emotional changes; 1–4) revealed no severe discomfort in any group ($F(2, 125) = 0.78$, $p = 0.46$, $\eta^2 = 0.01$).

#### 6. Delay-Invariance of tDCS Effects on Outcome and Aversiveness

Because the experimental paradigm manipulated time delays across trials, we examined whether the effects of tDCS varied as a function of temporal delay (i.e., immediate versus delayed outcomes; **Figure S11**). To test this, we conducted repeated-measures ANOVAs including within-trial delay as a within-subject factor and stimulation group as a between-subjects factor.

For sustainable decisions, the stimulation effects were stable across all time points.

Specifically, outcome ratings exhibited a significant between-subjects main effect of group ($F(2{,}125) = 22.496,\ p < .001$), but no Group $\times$Delay interaction ($F(8{,}500) = 0.470$, $p = .877$; Figure S10A) and no main effect of delay ($F(4{,}500) = 1.346,\ p = .252$). This indicates that group differences were delay-invariant and already evident at the shortest delays. Aversiveness ratings for sustainable options mirrored this pattern: a significant main effect of group ($F(2{,}125) = 8.154,\ p < .001$), but neither a Group $\times$Delay interaction ($F(8{,}500) = 0.776,\ p = .624$; Figure S10B) nor a main effect of delay ($F(4{,}500) = 1.276$, $p = .278$).

For the unsustainable context, outcome ratings similarly showed a significant main effect of group ($F(2{,}125) = 10.263,\ p < .001$) without a Group $\times$Delay interaction ($F(8{,}500) = 0.651,\ p = .734$; Figure S10C) or a main effect of delay ($F(4{,}500) = 2.100$, $p = .080$). In contrast, aversiveness ratings for unsustainable options showed no significant effects: the main effect of group ($F(2{,}125) = 0.600,\ p = .551$), the main effect of delay ($F(4{,}500) = 1.665,\ p = .157$), and the interaction ($F(8{,}500) = 0.554,\ p = .816$; Figure S10D) were all non-significant. Together, these results confirm that the perceived aversiveness of unsustainable options remained stable across both groups and delays.

#### 7. Effects of tDCS on Computational Parameters (Model 3)

We next investigated whether tDCS altered the underlying computational parameters of Model 3 by comparing pre-to-post stimulation change scores (Day 3 minus Day 1; denoted as $\Delta a$, $\Delta b$, and $\Delta c$) across the three experimental groups.

For sustainable behaviors, one-way ANOVAs revealed no significant group differences in any parameter change scores: $\Delta a(F(2{,}125) = 0.315,\ p = .731,\ \eta^2 = .005,\ \omega^2 = -.011$; active-DLPFC: $-0.095 \pm 0.687$, active-Oz: $0.023 \pm 0.975$, sham-DLPFC: $0.013 \pm 0.542$), $\Delta b(F(2{,}125) = 0.806,\ p = .449,\ \eta^2 = .013,\ \omega^2 = -.003$; active-

DLPFC: $0.123 \pm 0.462$, active-Oz: $0.010 \pm 0.373$, sham-DLPFC: $0.070 \pm 0.398$), and $\Delta c$($F(2,125) = 0.407$, $p = .666$, $\eta^2 = .006$, $\omega^2 = -.009$; active-DLPFC: $8.269 \pm 57.240$, active-Oz: $-0.975 \pm 77.542$, sham-DLPFC: $-3.449 \pm 50.269$).

For unsustainable behaviors, group differences for $\Delta a$ and $\Delta b$ did not reach the $\alpha = 0.05$significance threshold ($\Delta a$: $F(2,125) = 2.919$, $p = .058$, $\eta^2 = .045$, $\omega^2 = .029$; $\Delta b$: $F(2,125) = 2.719$, $p = .070$, $\eta^2 = .042$, $\omega^2 = .026$). Conversely, $\Delta c$exhibited a significant main effect of group ($F(2,125) = 5.632$, $p = .005$, $\eta^2 = .083$, $\omega^2 = .067$; active-DLPFC: $-10.705 \pm 37.967$, active-Oz: $15.093 \pm 33.885$, sham-DLPFC: $6.124 \pm 37.116$). Follow-up uncorrected Welch's t-tests confirmed that the active-DLPFC group differed significantly from both the active-Oz group (mean difference $= 25.797$, 95% CI $[10.441, 41.153]$, $p = .001$) and the sham-DLPFC group (mean difference $= 16.828$, 95% CI $[0.530, 33.127]$, $p = .043$). The active-Oz and sham-DLPFC control groups did not differ from one another (mean difference $= -8.969$, 95% CI $[-24.342, 6.404]$, $p = .249$).

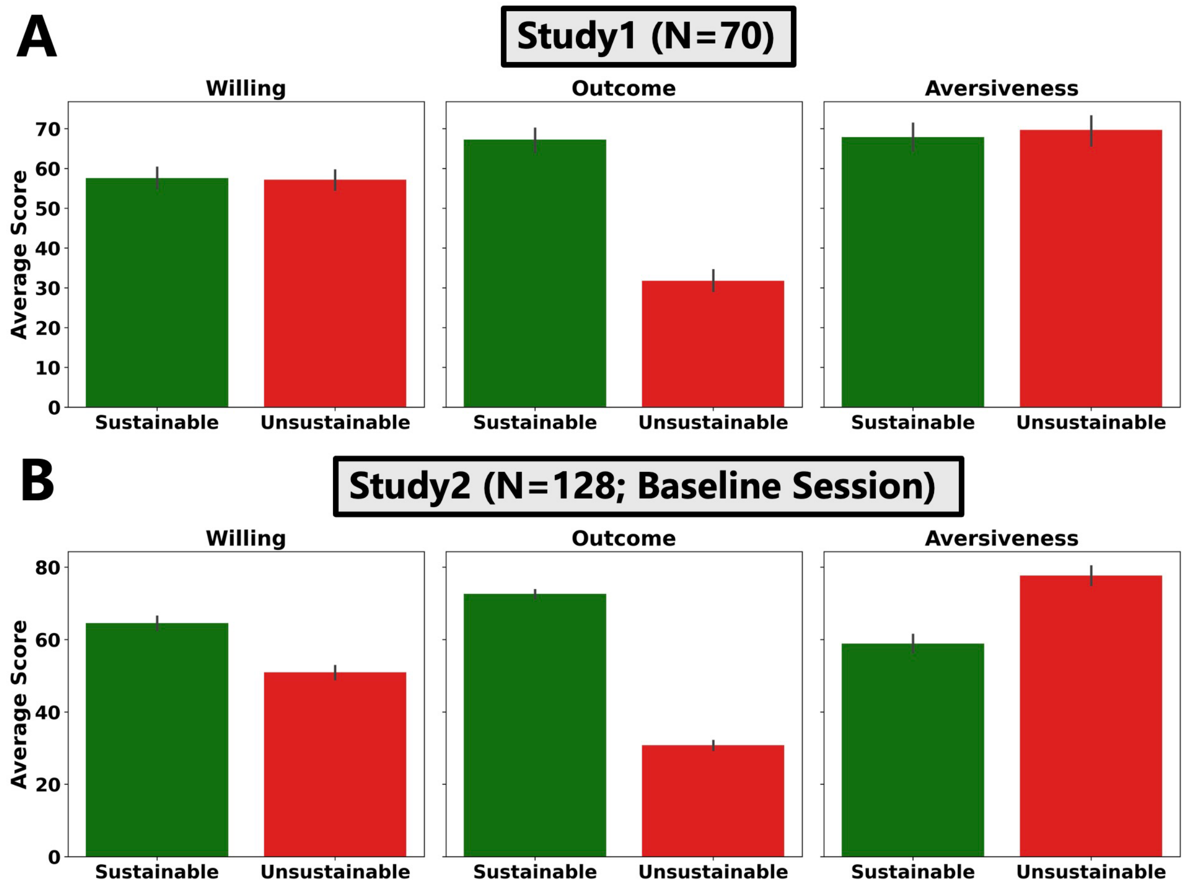


**Figure S1 Mean willingness, perceived outcome value, and perceived aversiveness for sustainable vs. unsustainable behaviors (Study 1 and Study 2 baseline)** (A) Study 1 (N = 70). (B) Study 2 baseline session (N = 128). For each participant, trial-level ratings were aggregated to obtain mean willingness to perform the behavior, perceived outcome value, and perceived task aversiveness

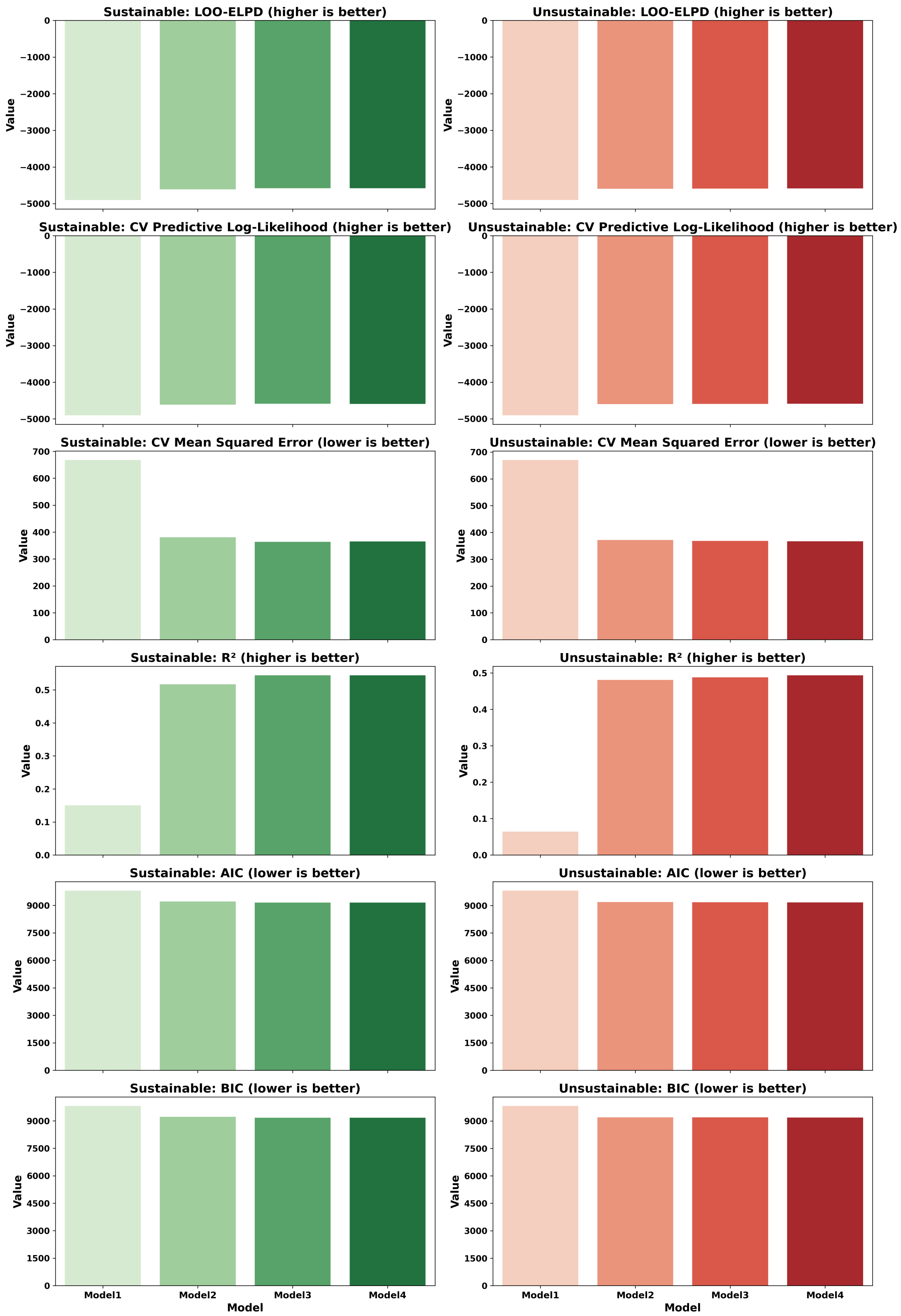


**Figure S2 Model comparison for predicting willingness from outcome value and aversiveness in Study 1 (sustainable vs. unsustainable trials).** Trial-wise willingness (y) was modeled as a function of perceived outcome value (O) and task aversiveness (A) using four nested Gaussian linear models estimated via OLS. Model performance was evaluated with (i) leave-one-out cross-validation expected log predictive density (LOO-ELPD; higher is better); (ii) AIC and BIC derived from the Gaussian log-likelihood; and (iii) participant-wise generalization via {5-fold GroupKFold cross-validation, keeping all trials from a participant in the same fold. CV metrics include predictive log-likelihood and mean squared error on held-out trials.

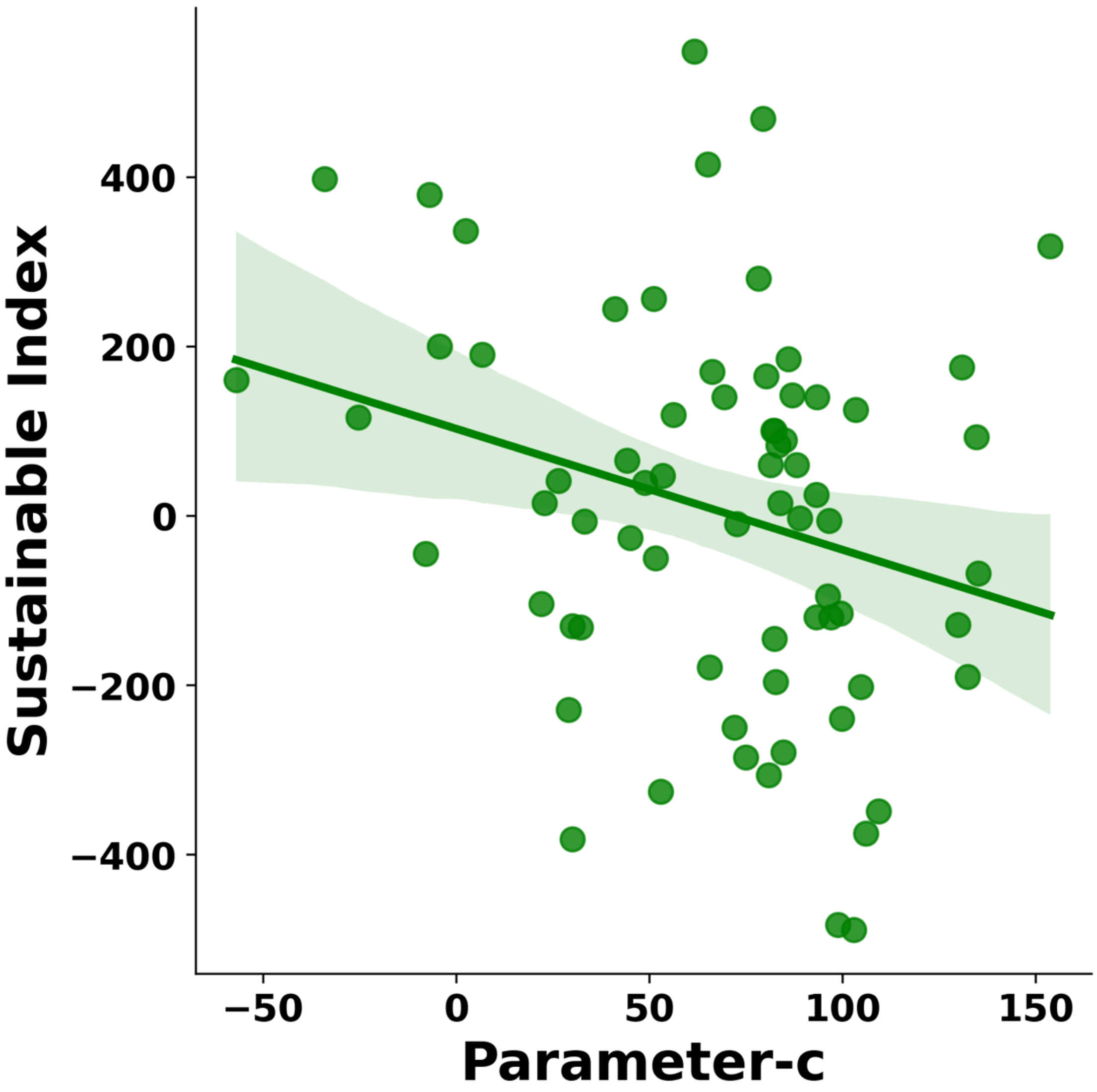


**Figure S3 Decision/bias parameter (c) estimated on sustainable trials was negatively associated with the sustainability index in Study1.**

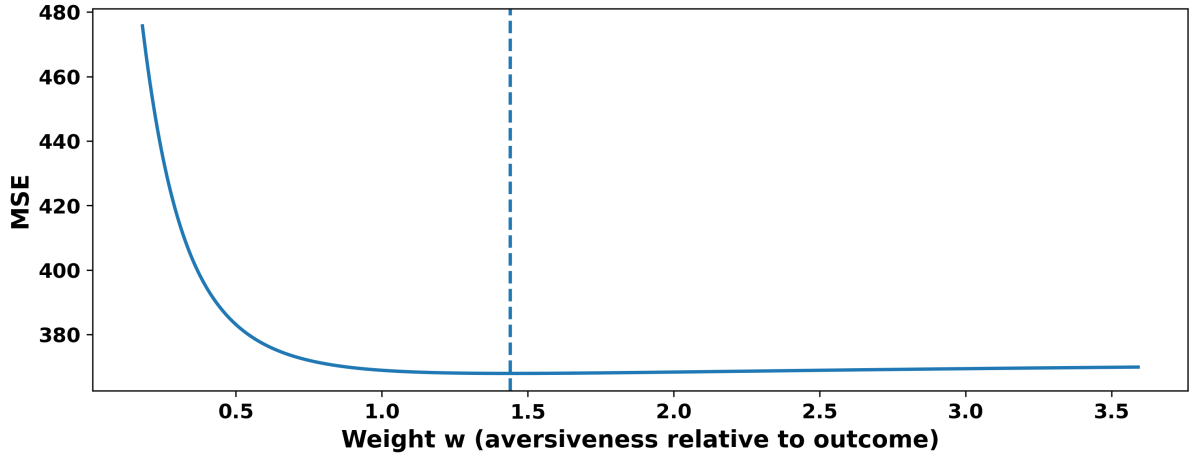


**Figure S4 Selecting the aversiveness weight (w) that best predicts willingness from a composite value signal.** For candidate weights (w) in (0-4) (step =0.001), we constructed a composite predictor [$x(w) = \text{Outcome} - w \cdot \text{Aversiveness}$] and fit [$\text{Willingness} = \alpha + \beta x(w) + \varepsilon$] using Study1. The curve shows MSE as a function of (w); the dashed line marks the selected optimum (w = 1.44).

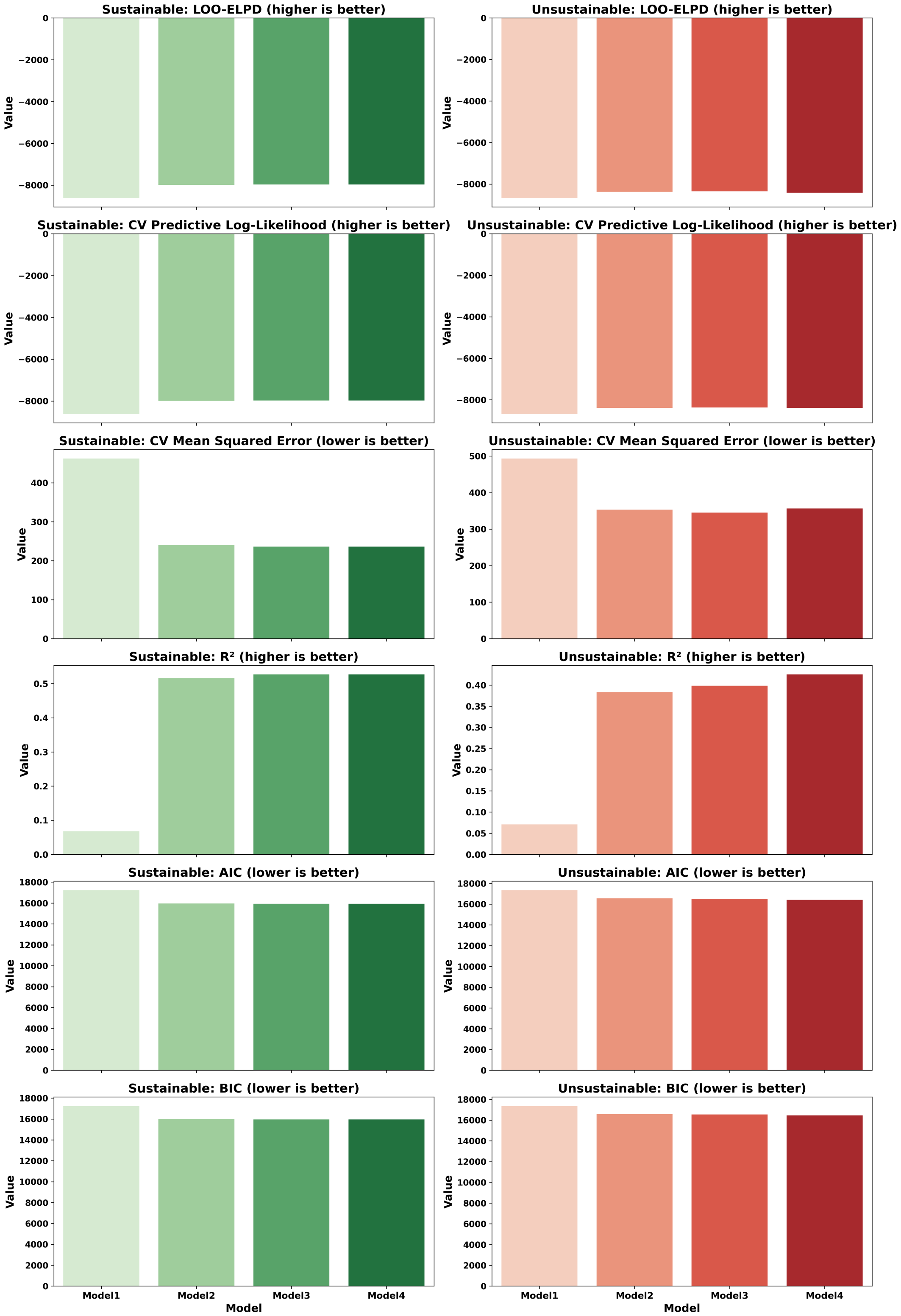


**Figure S5 Model comparison for predicting willingness from outcome value and aversiveness in Study 2 (sustainable vs. unsustainable trials).** Trial-wise willingness (y) was modeled as a function of perceived outcome value (O) and task aversiveness (A) using four nested Gaussian linear models estimated via OLS. Model performance was evaluated with (i) leave-one-out cross-validation expected log predictive density (LOO-ELPD; higher is better); (ii) AIC and BIC derived from the Gaussian log-likelihood; and (iii) participant-wise generalization via {5-fold GroupKFold cross-validation, keeping all trials from a participant in the same fold. CV metrics include predictive log-likelihood and mean squared error on held-out trials.

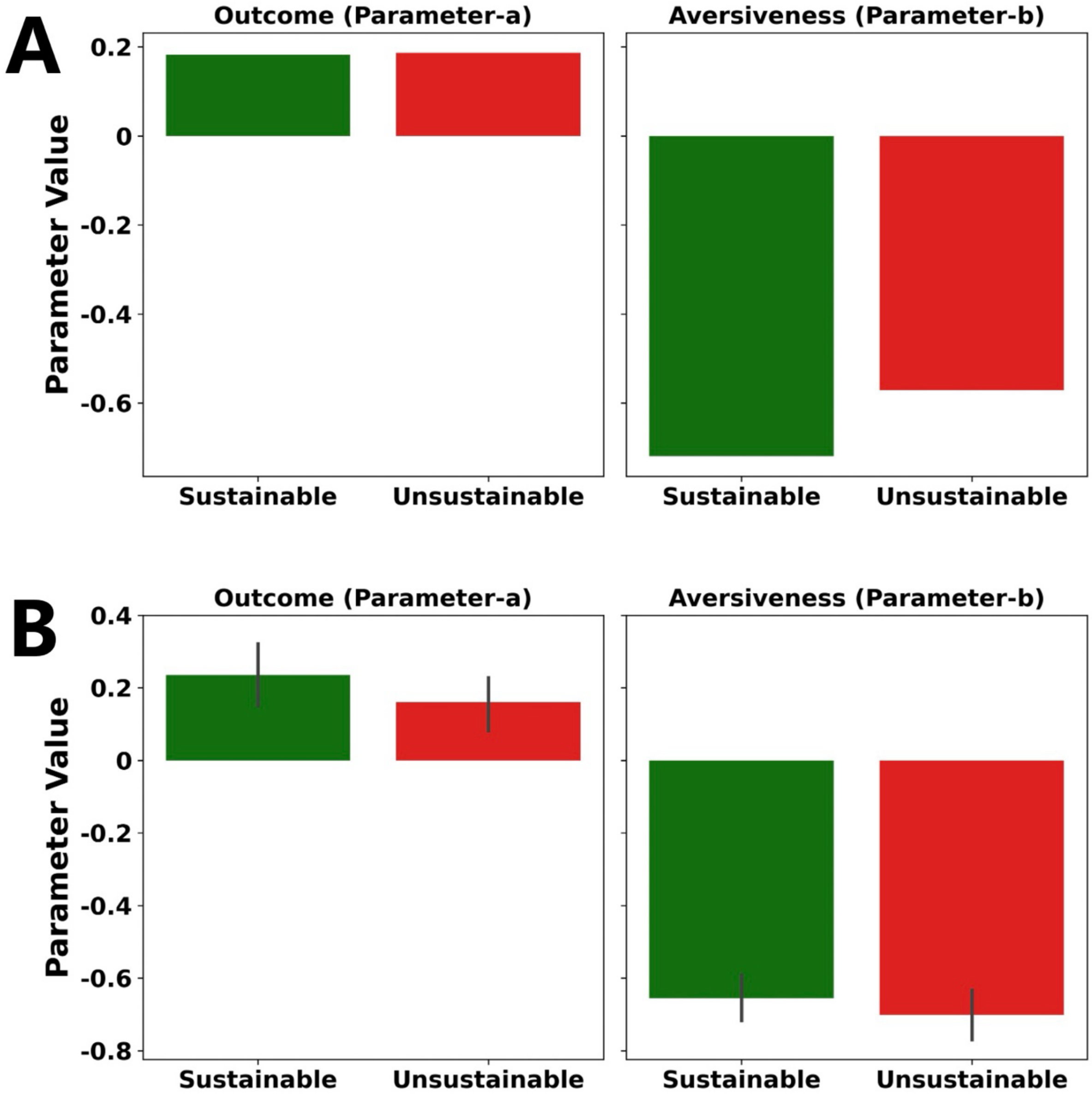


**Figure S6 Comparing computational sensitivities to outcome and aversiveness across sustainable and unsustainable contexts (Study 2).** Parameters were estimated f by fitting Model3 separately to sustainable and unsustainable trials. (A) Group-level model fitting. (B) Individual-level model fitting.

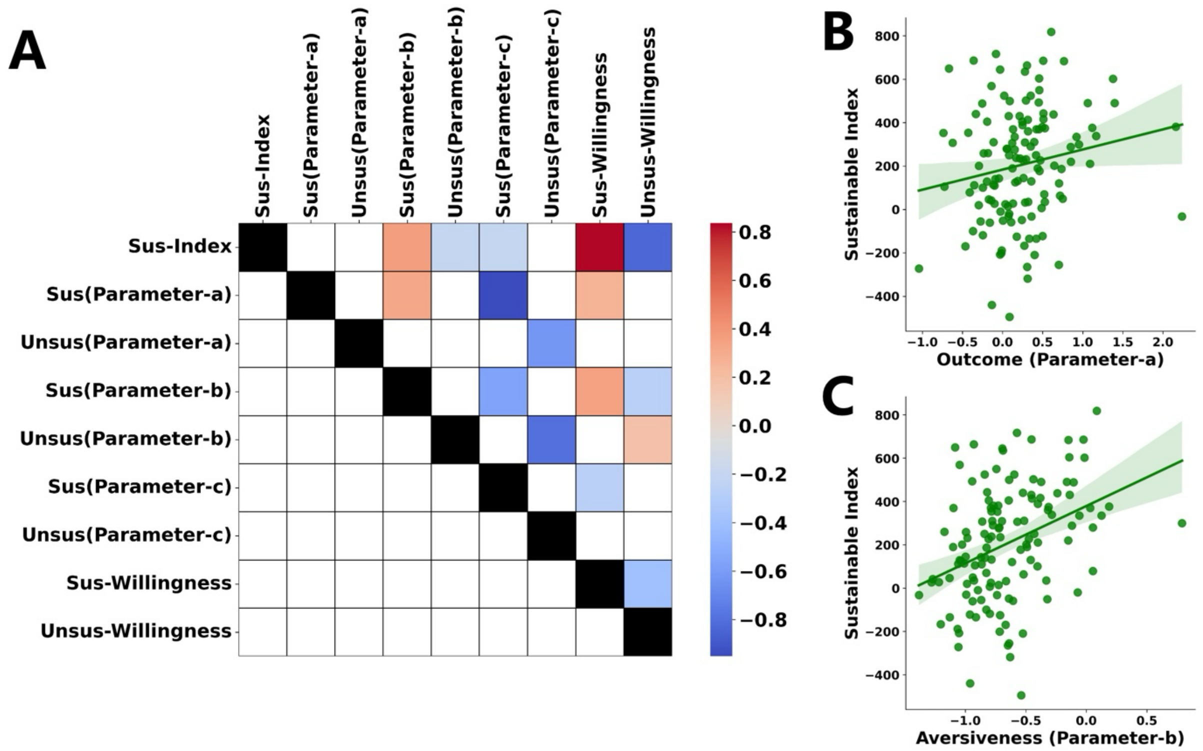


**Figure S7 Correlations between sustainability index, willingness, and computational parameters (a, b, c) in Study2.** We computed participant-level parameters (a, b, c) by fitting Model3 separately to sustainable and unsustainable trials, alongside mean willingness ratings in each context. (A) Correlation matrix including the sustainability index (Sus-Index), willingness ratings, and participant-level model parameters (a, b, c) estimated separately from sustainable (Sus) and unsustainable (Unsus) trials. (B–C) Scatter plots showing associations between the sustainability index and (B) outcome sensitivity (a) and (C) aversiveness sensitivity (b) from the sustainable-trial model.

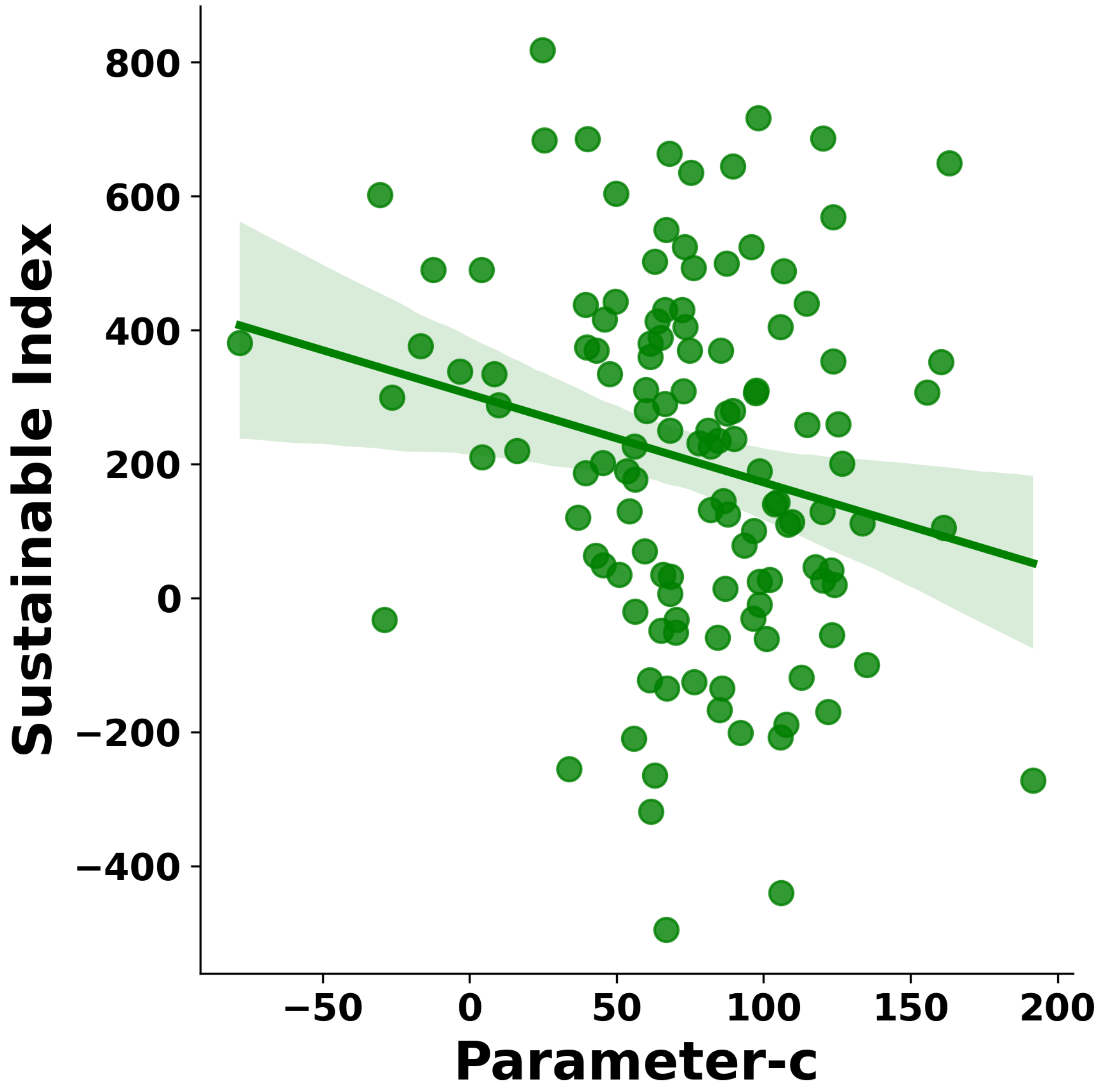


**Figure S8 Association between the sustainable-trial bias parameter (c) and the sustainability index (Study 2)**

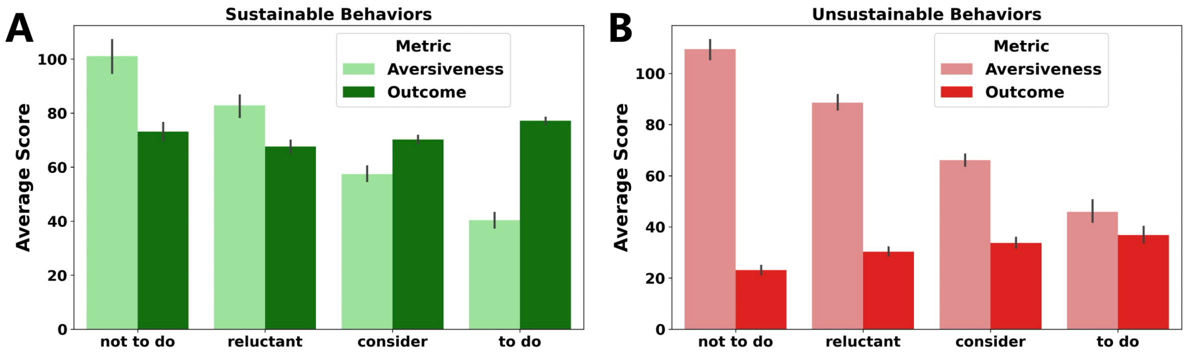


**Figure S9 Outcome–aversiveness balance across willingness-based decision categories for sustainable and unsustainable behaviors (Study 2).** Trials were stratified into four categories based on willingness ratings: "not to do," "reluctant," "consider," and "to do" (A) Individuals choose against sustainable behaviors when task aversiveness exerts a stronger influence than outcomes, but they choose to engage in behaviors when aversiveness is less influential than outcomes. (B) Outcome-aversiveness comparisons do not affect decisions regarding unsustainable behaviors, despite a significant decrease in aversiveness from the option not-to-do to option to-do.

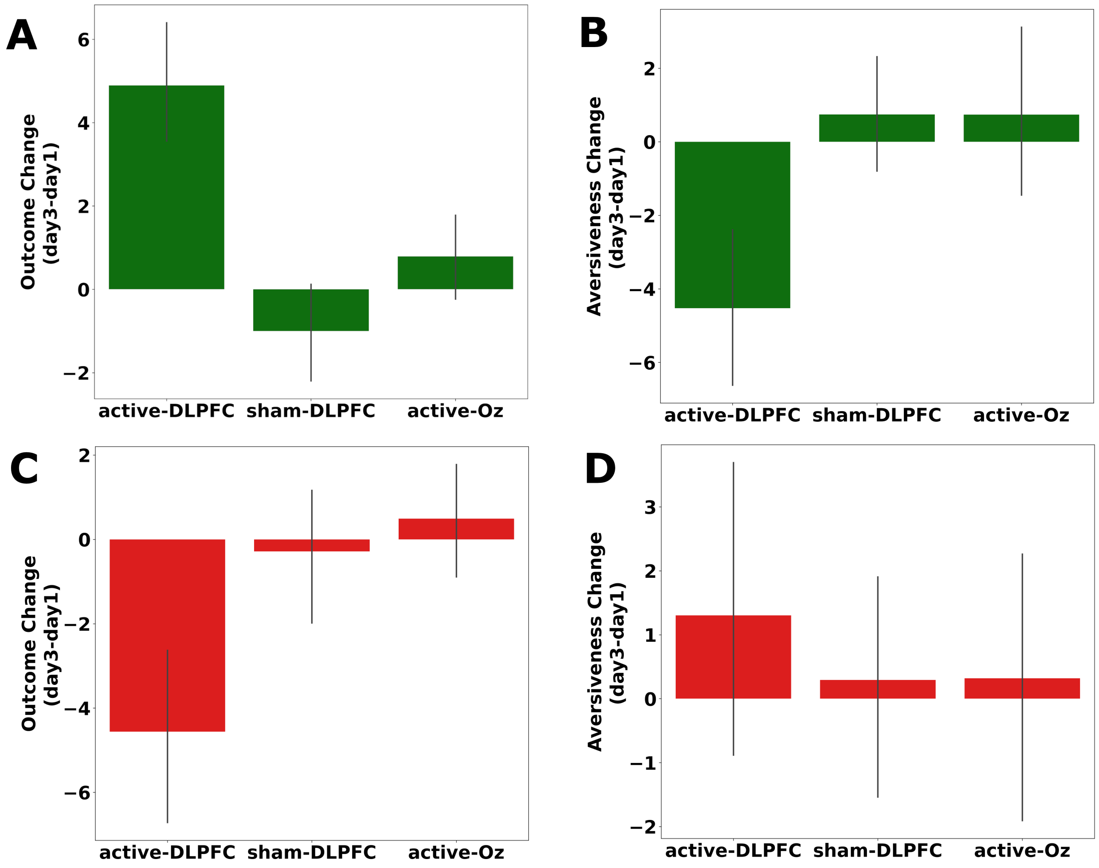


**Figure S10 tDCS group differences in change scores for evaluative components (outcome and aversiveness) in Study 2.** Stimulation-dependent changes in outcome and aversiveness ratings (Study 2). Change scores were computed as [$\Delta_{rating} = rating_{Day3} - rating_{Day1}$] for each participant and separately for sustainable and unsustainable trials. Participants were assigned to active-lDLPFC, sham-lDLPFC and active-Oz group . Panels A–D depict ΔOutcome and ΔAversiveness for sustainable and unsustainable trials (panel mapping specified explicitly).

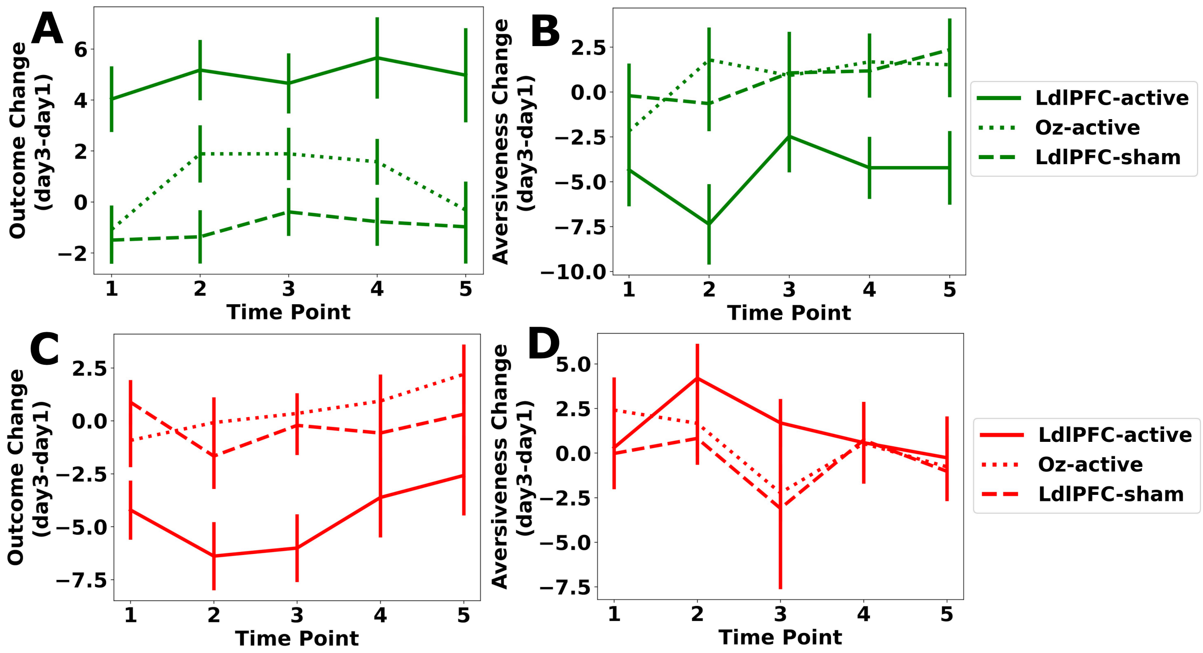


**Figure S11 Delay-invariance test of tDCS effects on outcome value and aversiveness (Study 2).** For each delay level (Time Points 1–5), we computed participant-level change scores as [$\Delta$ = Day3 − Day1] for outcome value and aversiveness ratings, separately for sustainable and unsustainable trials. Group trajectories are shown for active-lDLPFC, sham-lDLPFC, active-Oz separately.

### Supplemental Tables

**Table S1 Online evaluation results for the stimulus pool and the final selected subset (participant-rated; image-level summary) (n=480)**

| Score | Selected-Sustainable (n=16) | Selected-Non sustainable (n=16) | Unelected-Sustainable (n=20) | Unelected -Non sustainable (n=20) |
|---|---|---|---|---|
| **Q1-4 Total** | 23.55±1.06 | 23.70±1.82 | 23.62±0.92 | 22.09±2.14 |
| **Q4** | 5.80±0.21 | 6.01±0.28 | 5.61±0.26 | 5.51±0.33 |
| **Q5** | 5.99±0.48 | 3.64±0.51 | 5.32±0.48 | 4.22±0.59 |

**Note:** Participants rated 72 images on five 7-point Likert items (1–7). Q1–Q4 were summed to form a comprehensibility index (range 4–28). Q4 indexed ease of classifying environmental friendliness; Q5 indexed perceived environmental friendliness. Columns report image-set summaries for selected versus unselected images in sustainable and unsustainable categories. Selected images were provided in the OSF project folder (see methods, Data and Code availability).

**Table S2: Self-reported affect, perceived stimulation intensity, and adverse effects by stimulation group (Study 2)**

| tDCS Adverse Effect Assessment | | Active LDLPFC (n=43) | Sham LDLPFC (n=41) | Active Oz (n=44) |
|---|---|---|---|---|
| Emotion | Before stimulation | 11.05±3.56 | 10.81±3.14 | 11.00±2.20 |
| | After stimulation | 11.54±3.31 | 10.22±3.42 | 11.96±3.10 |
| Stimulation Intensity | | 5.20±1.75 | 4.15±1.91 | 4.89±1.93 |
| Discomfort | | 14.51±4.58 | 13.71±3.76 | 14.78±3.85 |

**Note:** Emotion was assessed before and after brain stimulation. Perceived stimulation intensity was rated on a 1–9 scale (1 = not intense at all, 9 = extremely intense). Discomfort reflects sum of adverse-effect items rated 1–4 (1 = none, 4 = severe).

### Supplemental Questionnaires

**Questionnaire to measure self-reported emotion, stimulation intensity, and discomfort following tDCS**

**Emotion self-report (1–9 scale)**
Please rate your current emotional state by selecting one number on each 1–9 scale shown below (1 = lowest, 9 = highest) and mark your response.

2. Current pleasantness

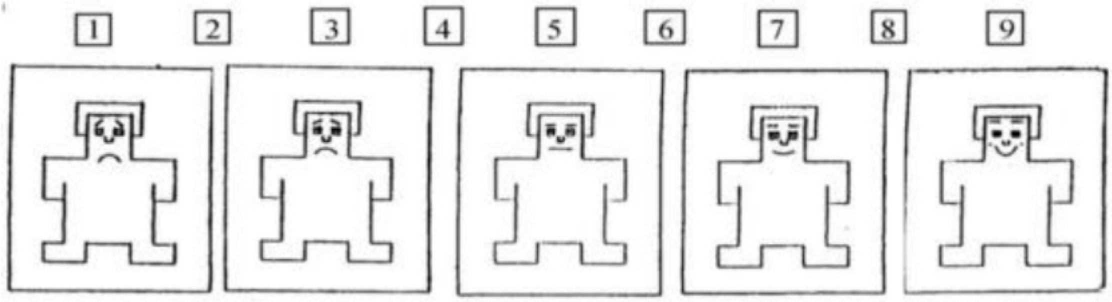


3. Current activation/arousal

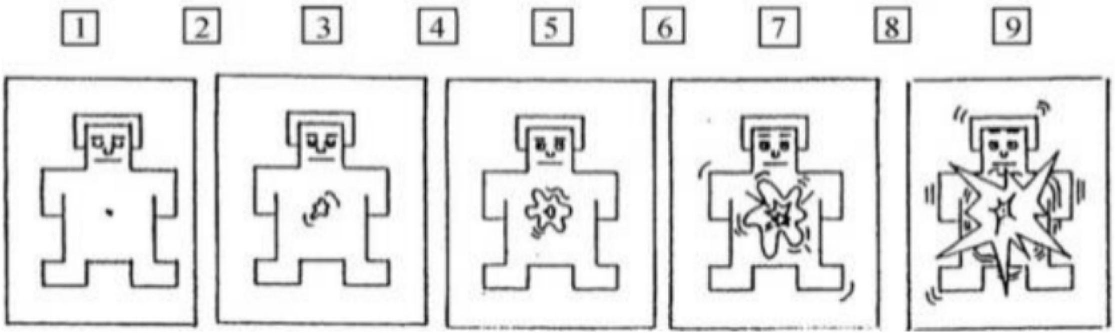


**Perceived stimulation intensity (1–9 scale)**
How intense do you think the stimulation you just received was? (1–9; 1 = not intense at all, 9 = extremely intense)

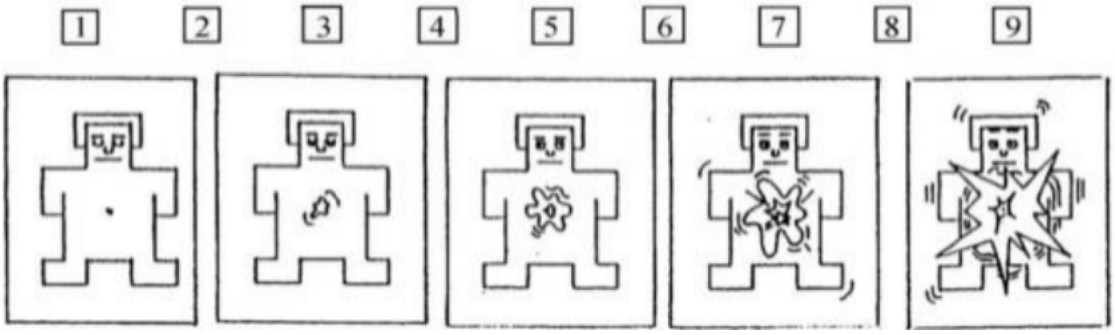

**Potential tDCS adverse reactions (1–4 scale)**

During the experiment/stimulation, did you experience any of the following sensations? Please rate each item on a 4-point scale:

| | none | mild | moderate | severe |
|---|---|---|---|---|
| Headache | 1 | 2 | 3 | 4 |
| Neck pain | 1 | 2 | 3 | 4 |
| Scalp pain | 1 | 2 | 3 | 4 |
| Numbness | 1 | 2 | 3 | 4 |
| Itching | 1 | 2 | 3 | 4 |
| Burning sensation | 1 | 2 | 3 | 4 |
| Skin redness | 1 | 2 | 3 | 4 |
| Sleepiness/drowsiness | 1 | 2 | 3 | 4 |
| Difficulty concentrating | 1 | 2 | 3 | 4 |
| Acute mood change | 1 | 2 | 3 | 4 |
| Other (please specify)<br>Severity (if rated) | 1 | 2 | 3 | 4 |

**The General Ecological Behavior (GEB) questionnaire**

Please indicate how often you engage in the following behaviors in your daily life:

| | never | rarely | sometimes | often | always |
|---|---|---|---|---|---|
| 1. I own energy efficient household devices | 1 | 2 | 3 | 4 | 5 |
| 2.I wait until I have a full load before doing my laundry | 1 | 2 | 3 | 4 | 5 |
| 3. I wash dirty clothes without prewashing | 1 | 2 | 3 | 4 | 5 |
| 4. In hotels, I have the towels changed daily | 1 | 2 | 3 | 4 | 5 |
| 5. I use a clothes dryer | 1 | 2 | 3 | 4 | 5 |
| 6. I bought solar panels to produce energy | 1 | 2 | 3 | 4 | 5 |
| 7. I use renewable energy sources | 1 | 2 | 3 | 4 | 5 |
| 8. In the winter, I keep the heat on so that I do not have to wear a sweater | 1 | 2 | 3 | 4 | 5 |
| 9. In the winter, I leave the windows open for long periods of time to let in fresh air | 1 | 2 | 3 | 4 | 5 |
| 10. In winter, I turn down the heat when I leave my apartment for more than 4 h | 1 | 2 | 3 | 4 | 5 |
| 11. I prefer to shower rather than to take a bath | 1 | 2 | 3 | 4 | 5 |
| 12. I drive my car in or into the city | 1 | 2 | 3 | 4 | 5 |
| 13. I drive on freeways at speeds under 100 kph (= 62.5 mph) | 1 | 2 | 3 | 4 | 5 |
| 14. I keep the engine running while waiting in front of a railroad crossing or in a traffic jam | 1 | 2 | 3 | 4 | 5 |
| 15. At red traffic lights, I keep the engine running | 1 | 2 | 3 | 4 | 5 |
| 16. I drive to where I want to start my hikes | 1 | 2 | 3 | 4 | 5 |
| 17. I refrain from owning a car | 1 | 2 | 3 | 4 | 5 |
| 18. I am a member of a carpool | 1 | 2 | 3 | 4 | 5 |
| 19. I drive in such a way as to keep my fuel consumption as low as possible | 1 | 2 | 3 | 4 | 5 |
| 20. I own a fuel-efficient automobile (less than 7 l per 100 km; i.e., less than 3 gallons per 100 miles) | 1 | 2 | 3 | 4 | 5 |

| | | | | | |
|---|---|---|---|---|---|
| 21. For longer journeys (more than 6 h), I take an airplane | 1 | 2 | 3 | 4 | 5 |
| 22. In nearby areas (around 30 km; around 20 miles), I use public transportation or ride a bike | 1 | 2 | 3 | 4 | 5 |
| 23. I ride a bicycle or take public transportation to work or school | 1 | 2 | 3 | 4 | 5 |
| 24. I buy milk in returnable bottles | 1 | 2 | 3 | 4 | 5 |
| 25. If I am offered a plastic bag in a store, I take it | 1 | 2 | 3 | 4 | 5 |
| 26. I reuse my shopping bags | 1 | 2 | 3 | 4 | 5 |
| 27. I buy beverages in cans | 1 | 2 | 3 | 4 | 5 |
| 28. I buy products in refillable packages | 1 | 2 | 3 | 4 | 5 |
| 29. I use fabric softener with my laundry | 1 | 2 | 3 | 4 | 5 |
| 30. I use an oven cleaning spray to clean my oven | 1 | 2 | 3 | 4 | 5 |
| 31. I kill insects with a chemical insecticide | 1 | 2 | 3 | 4 | 5 |
| 32. I use a chemical air freshener in my bathroom | 1 | 2 | 3 | 4 | 5 |
| 33. I buy convenience foods | 1 | 2 | 3 | 4 | 5 |
| 34. I buy seasonal produce | 1 | 2 | 3 | 4 | 5 |
| 35. I buy bleached and colored toilet paper | 1 | 2 | 3 | 4 | 5 |
| 36. I buy meat and produce with eco-labels | 1 | 2 | 3 | 4 | 5 |
| 37. I buy domestically grown wooden furniture | 1 | 2 | 3 | 4 | 5 |
| 38. I collect and recycle used paper | 1 | 2 | 3 | 4 | 5 |
| 39. I bring empty bottles to a recycling bin | 1 | 2 | 3 | 4 | 5 |
| 40. I put dead batteries in the garbage | 1 | 2 | 3 | 4 | 5 |
| 41. After meals, I dispose of leftovers in the toilet | 1 | 2 | 3 | 4 | 5 |
| 42. After a picnic, I leave the place as clean as it was originally | 1 | 2 | 3 | 4 | 5 |
| 43. I am a member of an environmental organization | 1 | 2 | 3 | 4 | 5 |
| 44. I read about environmental issues | 1 | 2 | 3 | 4 | 5 |
| 45. I contribute financially to environmental organizations | 1 | 2 | 3 | 4 | 5 |
| 46. I talk with friends about problems related to the environment | 1 | 2 | 3 | 4 | 5 |
| 47. I have pointed out unecological behavior to | 1 | 2 | 3 | 4 | 5 |

| | | | | | |
|---|---|---|---|---|---|
| someone | | | | | |
| 48. I boycott companies with an unecological background | 1 | 2 | 3 | 4 | 5 |
| 49. I have already looked into the pros and cons of having a private source of solar power | 1 | 2 | 3 | 4 | 5 |
| 50. I requested an estimate on having solar power installed | 1 | 2 | 3 | 4 | 5 |
